# Environmentally stressed human nucleus pulposus cells trigger the onset of discogenic low back pain

**DOI:** 10.1101/2023.03.02.530506

**Authors:** Wensen Jiang, Juliane D Glaeser, Giselle Kaneda, Julia Sheyn, Jacob T Wechsler, Stephen Stephan, Khosrowdad Salehi, Julie L. Chan, Wafa Tawackoli, Pablo Avalos, Christopher Johnson, Chloe Castaneda, Linda EA Kanim, Teerachat Tanasansomboon, Joshua Burda, Oksana Shelest, Haneen Yameen, Tiffany G Perry, Michael Kropf, Jason M Cuellar, Dror Seliktar, Hyun W Bae, Laura S Stone, Dmitriy Sheyn

## Abstract

Low back pain (LBP) is often associated with the degeneration of human intervertebral discs (IVDs). However, the pain-inducing mechanism in degenerating discs remains to be elucidated. Here, we identified a subtype of locally residing nucleus pulposus cells (NPCs), generated by the environmental stress in degenerating discs, that triggered the onset of discogenic LBP. Single-cell transcriptomic analysis of human tissues showed a strong correlation between this specific pain-triggering subtype and the pain conditions in human degenerated discs. Next, we recreated this pain-triggering subtype by applying known exogenous stressors to healthy NPCs *in vitro*. The recreated pain phenotype activated functional sensory neurons response *in vitro* and induced local inflammatory responses, hyperalgesia, and mechanical sensitivity in a healthy rat IVD *in vivo*. Our findings provide strong evidence of a previously unknown pain-inducing mechanism mediated by NPCs in degenerating IVDs. This newly defined pathway will aid in the development of NPC-targeted therapeutic strategies for clinically unmet need to attenuate discogenic LBP.

**One Sentence Summary:** Discogenic low back pain can be initiated by a stress-induced subtype of nucleus pulposus cells present in human degenerating intervertebral discs

## INTRODUCTION

Low back pain (LBP) is a leading cause of disability worldwide, affecting almost 80% of the adult population (*1, 2*). Approximately 40% of LBP cases are attributed to intervertebral disc (IYD) degeneration (*3, 4*). Previous studies have established that deterioration of the disc structure, caused by endogenous or exogenous triggers, leads to the vascularization of the inner region of the disc, the Nucleus Pulposus (NP) (*5–7*). Immune cells then migrate into the NP and secrete inflammatory cytokines that may lead to innervation of the IYD and pain (*5–7*). However, this theory does not explain the clinical observation that some degenerating IYDs do not exhibit discogenic pain phenotype (*8–15*). The underlying mechanism that determines which degenerating disc would drive discogenic LBP is yet to be revealed (*5, 16*).

Multiple environmental triggers are implicated in IYD degeneration (e.g., overloading, inflammation, and low pH) (*17–20*). Pro-inflammatory cytokines including IL-1B and TNF-a can induce IYD cell senescence, apoptosis, and matrix degradation (*21*). Due to the low supply of oxygen in the avascular NP, IYD cells undergo lactic acid fermentation at a higher rate, decreasing pH levels (*22*). The introduction of mechanical stress to cartilage, a tissue with similar properties to the NP, increases oxidant production and decreases the functionality of the cartilage tissue. These processes are all linked to the IYD degeneration (*23*). Although much has been studied regarding the etiology of IYD degeneration, it is still unclear which factors transform degenerating IYDs into painful ones.

The human IYD consists of a jelly-like, proteoglycan-rich NP in the inner core, surrounded by a ring-like annulus fibrosus (AF) in the outer region, with two cartilaginous endplates adjoining the vertebral bodies (*24, 25*). Healthy NP tissue is avascular and aneural (*5, 6, 26-29*), with IYD nerve ingrowth only occurring in the outer one-third of the AF (*5, 29*). Despite preliminary investigations, the exact role of nucleus pulposus cells (NPCs) in discogenic LBP induction is still unknown (*5–7, 30*). Environmental triggers were found to affect NPC gene expression (*31–34*). Richardson et al. suggested that stressed NPCs can induce neural response (*35*), but no study has provided sufficient evidence for pain-inducing mechanisms mediated directly by NPCs. Most previous studies focused on the role of NPCs in maintaining the extracellular matrix. The loss or malfunction of the NPC correlates with matrix degeneration in the IVD (*31, 36*). Some studies reported that replacing or replenishing resident NPCs can potentially regenerate IVDs and treat discogenic LBP (*37–39*), but these studies did not specifically differentiate between IVD regeneration and pain attenuation.

Single-cell RNA sequencing (scRNA-seq) technology can be used to determine cell subtypes, identify rare cell populations, and discover novel biological markers of IVD cells (*40–42*). ScRNA-seq technology could establish mechanisms of discogenic LBP specific to NPC sub-populations. For example, there was limited understanding of the divergent NPC subtypes (*33*) until single-cell technologies led to the discovery of heterogeneous populations of cells residing in the IVD (*40–42*). The present study investigates the role of NPC subtypes in mediating discogenic LBP by leveraging scRNA-seq of patient-derived cells and validates the findings in both *in vitro* and *in vivo* models of discogenic pain. We hypothesize that NPC subtypes that are associated with discogenic pain development could be identified. We further hypothesized that this cellular phenotype could be reproduced by environmental stressors *ex vivo*, and that these stressed cells could then trigger the onset of discogenic pain in non-degenerated IVD. We first validated our hypothesis by showing the association between the pain conditions of IVDs and the single-cell transcriptomes of NPCs in human patients and cadaveric donor samples and identified the specific pain-associated NPC subtypes. Next, we recreated such pain-associated subtypes of NPCs by applying environmental stressors *in vitro*. The recreated pain phenotype was used as an *in vitro* cell model to study the direct crosstalk between the NPCs and nociceptors. Lastly, a rat model was utilized to validate the pain-triggering effects of the stressed NPCs *in vivo*.

Pain is a sensory and emotional experience and as such cannot be reproduced *in vitro*. However, it is possible to study the interactions between NPCs and sensory neurons (nociceptors) in model systems. We therefore established a microfluidic chip that allowed for physically separated co-culturing of NPCs and nociceptors while maintaining the molecular crosstalk and axon ingrowth between the two cell types. We utilized induced pluripotent stem cells (iPSC) to generate iPSC-derived nociceptor-like cells (iNOCs) for *in vitro* studies.

The rat model of discogenic pain is widely used in the LBP field, due to our ability not only to quantitatively measure the structural properties of the IVD post-degeneration and post-treatment but also to use biobehavioral tests to measure the pain outcomes as closely as possible to the clinical observations. The stimulus-evoked mechanical and thermal hyperalgesia tests, including von Frey and hot plate tests are commonly used a surrogate of LBP (*43*). Induction of IVD degeneration and discogenic pain is usually modeled *in vivo* by the IVD injury (*44–47*). We have previously evaluated the degree of IVD degeneration and discogenic pain via µMRI, gene expression analysis, histology, and biobehavioral tests (*47*). Here we used a healthy IVD for cell injections and measured the pain outcomes.

In this study, specific cell subtypes mediating the transformation from asymptomatic to painful discs and their molecular signatures associated with the pain induction were determined, collectively validating our hypothesis. The unveiling of NPC-mediated pain mechanism will inform NPC-targeted therapeutic strategies focusing on the clinical problem of discogenic LBP.

## RESULTS

### 3.1 Single-cell analyses identified subtypes of NPCs associated with discogenic LBP

The research strategy and experiment design are shown in Fig. 1. ScRNA-seq was performed on four asymptomatic IVD samples (aIVD, n=4) and three back pain IVD samples (bpIVD, n=3) and combined to create an integrated cell atlas for all IVD cells (Fig. 2A). The unsupervised clustering identified 12 clusters (Fig. 2A). We used previously reported markers to assign cell identifies to clusters, i.e., COL2A1, ACAN, and SOX9 for Nucleus Pulposus cells (NPCs)(*48–51*), COL1A1 for annulus fibrosus cells (AFCs)(*52, 53*), CRTAC1 for fibro-chondrocytes (FCs)(*54*), CD14 and CD44 for immune cells (ICs), (*55, 56*) and HBB and HBA for red blood cells (RBCs, Fig. 2A)(*57, 58*). Seven clusters were assigned as NPC subtypes (NPC1-7), two clusters were assigned to AFC subtypes (AFC1 and AFC2), and FC, IC, and RBC populations were found to be represented by one cluster (Fig. 2A). The expression level and the percentage of cells expressing the markers in each cluster are shown in Fig. S3A. All NPC subtypes expressed classical NPC markers and lacked AFC marker (Fig. 2B). The NPC populations were further classified into subtypes using the *in silico* generated top variable markers for each subtype (Fig. 2C). The NPC1-7 subtypes were identified using the expressed top markers MMP3, SPP1, S100A2, CLIP, HSPA6, PRG4, and MT-ATP6 respectively (Fig. 2C).

**Figure 1.**
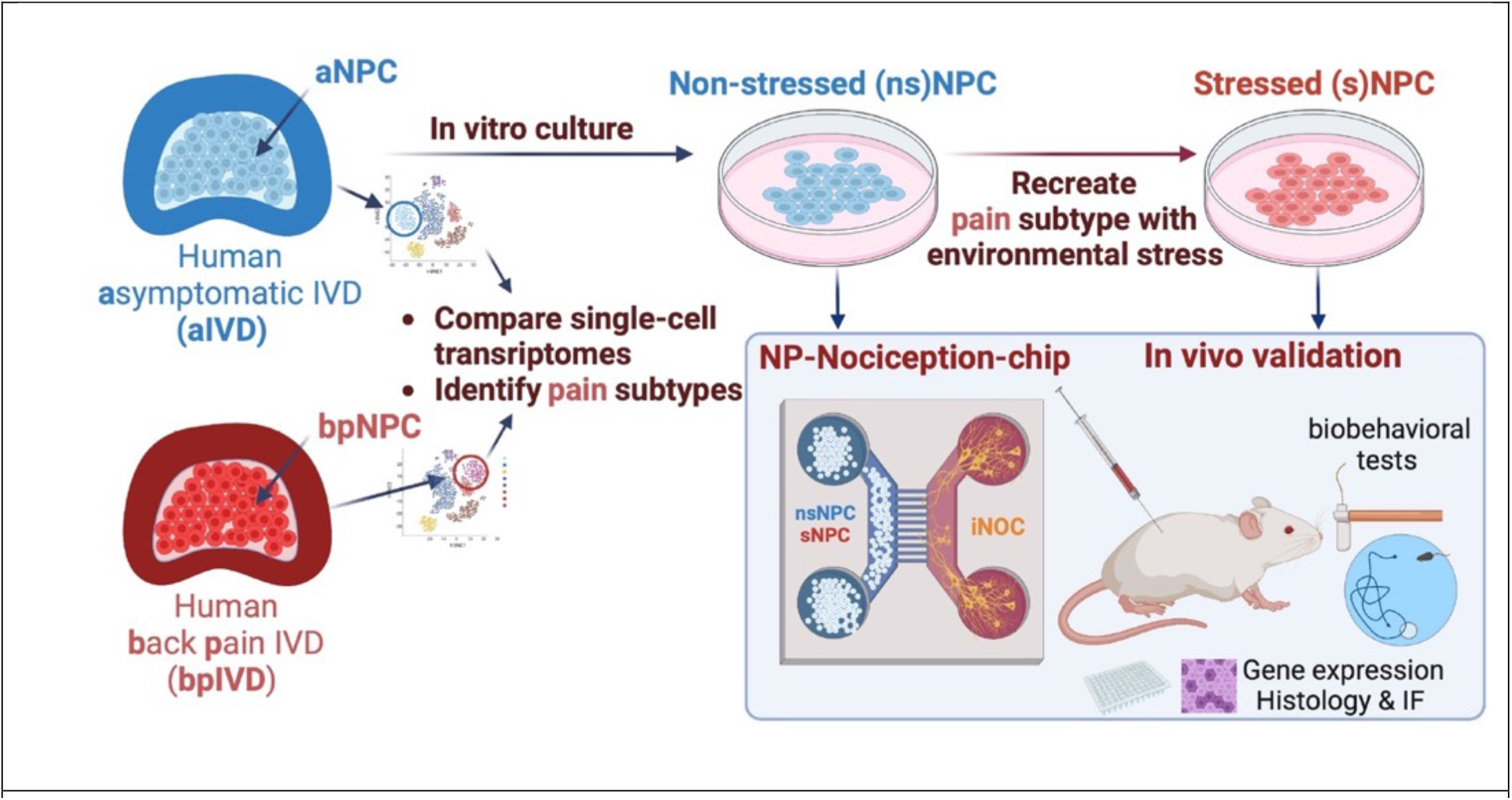
Illustrative scheme for research strategy and the experimental design.

**Fig. 2.**
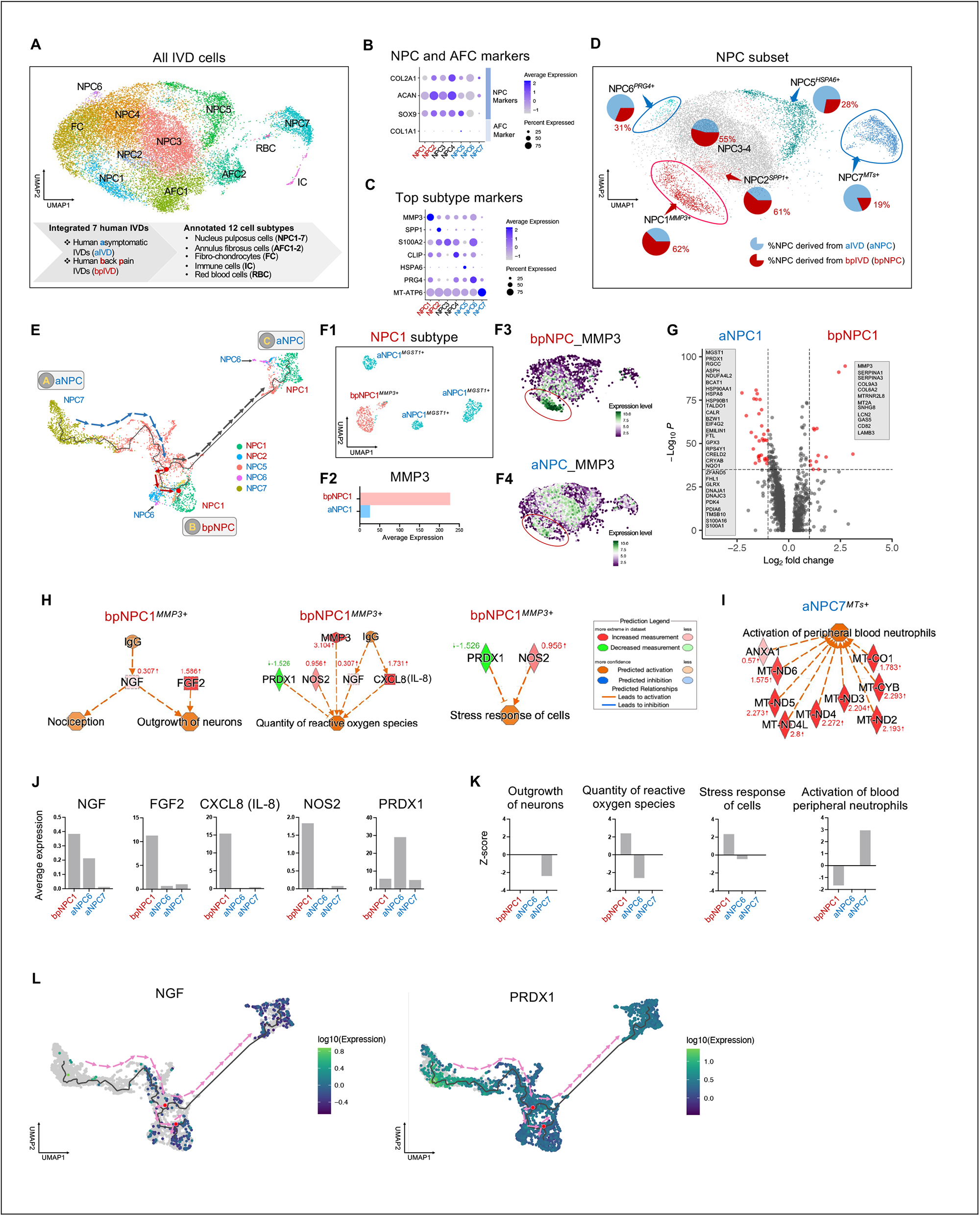
The heterogeneity of NPC populations and their transcriptomes in human IVDs is associated with LBP symptoms. **A)** Integrated single-cell atlas (UMAP) of IVDs from 7 patients. Dot plots of gene expression for markers labeling **B)** NPCs and AFCs and **C)** NPC subtypes. **D)** Subset atlas (UMAP) of NPCs with pie charts showing the percentage of cells in each cluster derived from different IVDs. Percent in red color shows cells derived from bpIVDs in the respective subtype. **E)** NPC subtypes with preferred IVD associations (NPC1, 2, 5, 6, and 7) were ordered on a pseudo-time developmental trajectory. **F1-F4)** NPCs have distinct transcriptomes despite being in the same cluster, manifested by **F1)** UMAP within NPC1, **F2)** average expression of NPC1 marker, MMP3, and the feature plots showing the MMP3 expression on the UMAP of NPC populations derived from **F3)** bpIVDs or **F4)** aIVDs. **G)** Volcano plot shows the differentially expressed genes in cluster NPC1 of cells derived from bpIVDs (bpNPC1) versus from aIVDs (aNPC1). Enriched networks of interest found in **H)** bpNPC1 and **I)** aNPC7. **J)** The expression levels of key regulators and **K)** enriched functions of relevant subtypes. **L)** NGF and PRDX1 expression projected on a pseudo-time trajectory.

We found an association between the NPC subtypes and the origin of the IVDs (Fig. 2D). The NPC populations consisted of seven subtypes with different prevalence to IVD origins (Fig. 2D). Specifically, we found that over 60% of NPC1-2 cell subtypes were derived from bpIVD, whereas over 60% of NPC5-7 cells were derived from aIVDs. NPCs derived from aIVDs were hereafter referred to as aNPCs, and NPCs derived from bpIVDs as bpNPCs.

Pseudo-time trajectory suggests a process of branched development (Fig. 2E). The trajectory originates from NPC7 (MTs+) at terminal A and then branches into two directions: bpNPC-dominated terminal B and aNPC-dominated terminal C, both consisted of NPC1 (MMP3+) and NPC6 (PRDX1+) subtypes. The arrows show the direction of pseudo-time (Fig. 2E). The color-coded trajectories demonstrating IVD origin or pseudo-time are shown in the supplemental information (Fig. S3B). This type of development of the pain associated NPC type from the asymptomatic IVD-associated NPCs, may indicate the effect of the degenerated environment of these cells, rather than genetical predisposition of certain cells from the beginning, although there is not enough evidence to exclude predisposition at this point.

NPC1 cells derived from different IVDs showed distinct transcriptome even though they belong to the same subtype (Fig. 2Fl-G). Therefore, our following analyses distinguished NPC1 cells derived from aIVDs or bpIVDs, hereafter referred to as aNPC1 or bpNPC1. The bpNPC1 cells showed much higher expression levels of MMP3 (Fig. 2F2-F4) and different top-expressed genes (Fig. 2G) than the aNPC1 cells. The top markers of aNPC6 and aNPC7 are also different from bpNPC6 and bpNPC7 respectively (Fig. S3C-D). The highly expressed genes of bpNPC1 cells were compared to other bpNPCs (Fig. S3E).

Several enriched networks of interest were identified for the bpNPC1 cells (Fig. 2H). In the first network, FGF2 and NGF expression leads to an increase in the “outgrowth of neurons” function, and NGF, regulated by IgG, positively affects “nociception”. In the second network, the expression of MMP3, NOS2, NGF, and CXCL8 (IL-8) increases the “quantity of reactive oxygen species (ROS)” function. The expression of NGF and CXCL8 (IL-8) is regulated by IgG. In this network, PRDXl, an inhibitor of the “quantity of reactive oxygen species” function, is downregulated. In the third network, the function “stress response of cells” is negatively regulated by PRDXl and positively regulated by NOS2. The downregulation of the inhibitor PRDXl and expression of NOS2 synergistically lead to the upregulation of the “stress response of cells” function. Another network of interest identified in the aNPC7 cells indicates that the expression of various mitochondrial genes (MTs, Fig. 2I) and ANXAl co-upregulated the “activation of peripheral blood neutrophils” function. The expression of key genes in these networks was quantified to elucidate the unique molecular signature of either bpNPC1 or the aNPC6-7 subtypes (Fig. 2J). The bpNPC1 cells showed much higher expression levels of NGF, FGF2, IL-8, and NOS2 regulators compared to that of aNPC6 and aNPC7 cells. However, they showed a lower expression of PRDXl than aNPC6 cells. Among aNPC cells, aNPC6 cells showed much higher NGF and PRDXl expression than aNPC7 cells but lower FGF expression than aNPC7 cells. We found that HIF-la is expressed in both bpNPCs and aNPCs but in different areas on the UMAP: in NPC3-NPC4 area in bpNPCs and in NPC7 area in aNPCs (Fig. S3F). We also found that both BNIP3 (Fig. S3G) and VEGF-A (Fig. S3H) are expressed only in NPC5 and NPC7 area in aNPCs, and not expressed in bpNPCs. SEMA3A and CRCP were both found with higher expression levels in aNPC7 than aNPC6 and bpNPC1 (Fig. S3I). When comparing between the entire bpNPC and aNPC populations, NGF, IL-8, was higher in bpNPCs (Fig. S3J). The NGF level expressed in bpNPC1 was higher than in total bpNPCs (Fig. S3J). However, we found extremely low expression of BDNF and TNF when compared to NGF (Fig. S3J).

Z-scores of enriched biological functions in these networks were identified (Fig. 2K). The bpNPC1 cells showed upregulated “quantity of reactive oxygen species” and “stress response of cells” and downregulated “activation of peripheral blood neutrophils” functions (Fig. 2I). The aNPC6 cells showed downregulated “quantity of reactive oxygen species” and “stress response of cells” (Fig. 2I). The aNPC7 cells showed downregulated “outgrowth of neurons” and upregulated “activation of peripheral blood neutrophils” (Fig. 2I).

We found that NGF and PRDXl expression correlates with pseudo-time (Fig. 2L). NGF showed an increased expression level along the pseudo-time on the trajectory, but PRDXl oppositely demonstrated a decreased one (Fig. 2L).

### 3.2 Environmentally stressed NPCs express inflammatory and degenerative markers *in vitro*

The viability of NPCs *in vitro* was significantly reduced at low pH (6.5) after 4-day culture compared to higher pH levels (Fig. 3C). Interestingly, statistically higher cell viability was detected when the media was modified to pH=7.5 compared to the control media (pH=7.84). The viability of NPCs was significantly reduced when cyclic mechanical compression at 22.2% and 30% strain was applied on 30 NPC-loaded hydrogel constructs, compared to the no mechanical compression group (0% strain). No significant change in NPC viability was found in the application of IL-1β 0-5 ng/ml) for 1 day or, low glucose media (0.1-3.1 g/ml) for up to 14 days.

**Fig. 3.**
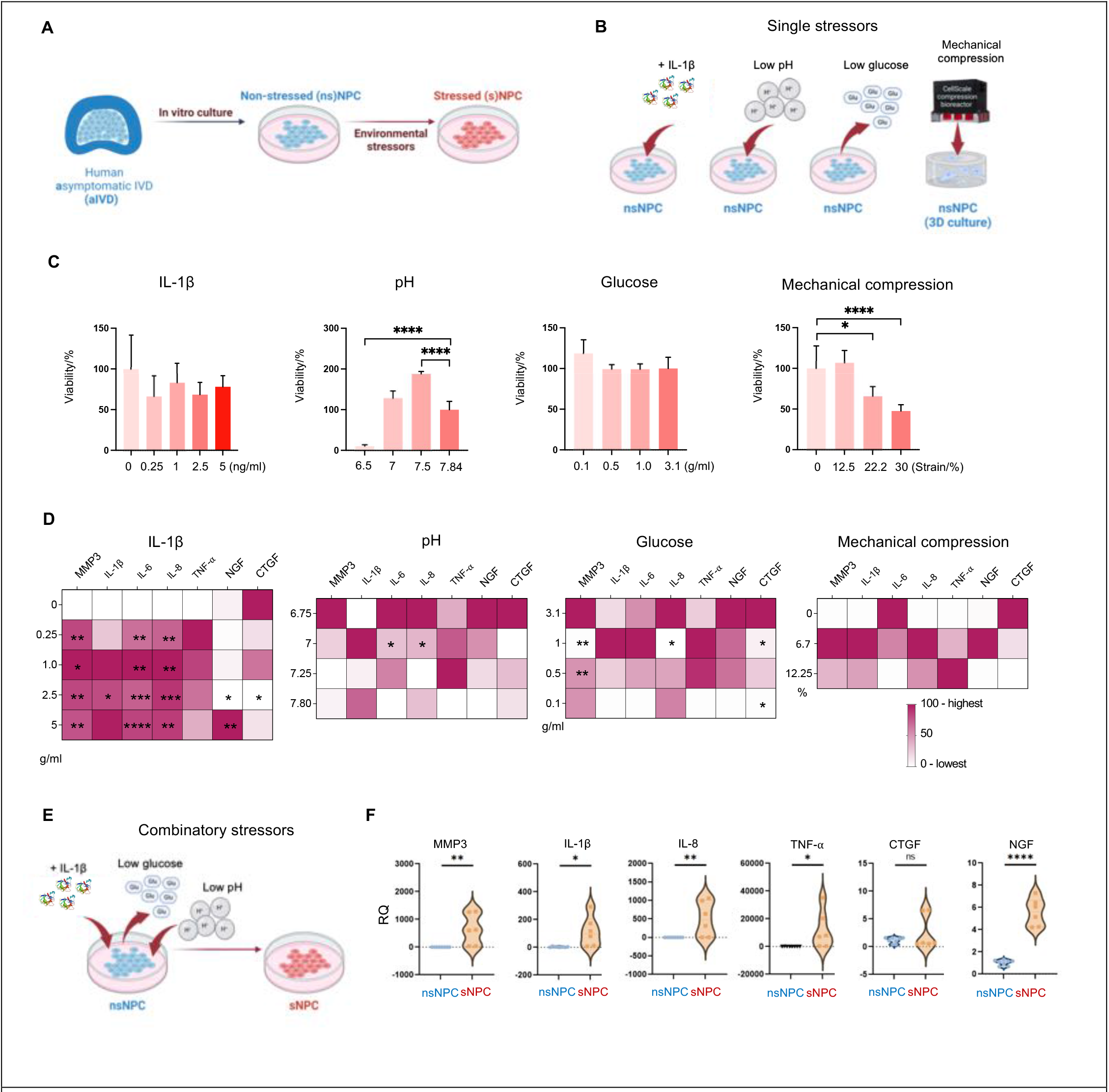
sNPC generated *in vitro* expressed inflammatory and degenerative markers. The experimental design for **A)** the *in vitro* model and **B)** single stressor screening. **C)** NPC viability under single stressors including inflammatory cytokines (IL-1β), low pH, low glucose level, and mechanical compression (n=4). **D)** Heatmap showing the expression of inflammatory and degenerative markers under single stressors (n=4), normalized from 0 (lowest) to 100 (highest) for each marker. **E)** The experimental design concept for the implementation of combinatory stressors applied to NPCs. **F)** The expression levels of inflammatory and degenerative markers under combinatory stressors (n=6) in terms of relative quantification (RQ). The nsNPCs control group for **F)** were cultured in unmodified media (pH=7.80-7.84, 3.1 g/ml glucose), no mechanical compression or IL-1β addition. (\**p*<0.05, \*\**p*<0.01, \*\*\**p*<0.001, \*\*\*\**p*<0.0001, ns or not labeled *p*>0.05)

NPCs exposed to different levels of IL-1β for 1 day highly expressed MMP3, IL-6, IL-8, and IL-1β at statistically higher levels than the control group (Fig. 3C). Interestingly, IL-1β at 2.5ng/ml downregulated NGF and CTGF levels, but at 5ng/ml the expression of NGF was upregulated. The pH7.25 induced IL-6 expression but suppressed IL-8 expression compared to the control group (pH=7.8). The pH7.0 and pH6.75 appeared to upregulate the expression levels of various markers, but no statistical difference was detected for any gene compared to the control. The glucose at the physiological level (1g/ml) induced NPCs to express MMP3 but suppressed the expression of IL-8 and CTGF with statistical significance compared to the control (Fig. 30). Further decreasing glucose levels to 0.5g/ml and 0.1g/ml downregulated MMP3 and CTGF genes respectively with the statistical difference compared to the control group (Fig. 30). Lastly, mechanical compression did not change the expression levels of any marker with statistical significance, but it appears to increase the expression of MMP3, IL-1β, IL-8, TNF-α, and NGF at 6.7% compression strain.

The chosen combined stressors aimed to maximize the stressing effects on NPCs while maintaining sufficient cell viability (Fig. 3E). We chose IL-lβ (2.5 g/ml), pH (6.75), and glucose (l g/ml) in 20 culture as the conditions of combined stressors. The mechanical compression stressor was not included in the combined stressors. In the range that the viability was not reduced (strain 0-l2.5%, Fig. 3C), the mechanical compression did not demonstrate statistically significant expression of stress markers (Fig. 30). Moreover, the 30-loaded NPCs are difficult to extract from the hydrogel construct for the following studies.

The combinatorically stressed NPCs (sNPC) highly expressed inflammatory and degenerative markers MMP3, IL-lβ, IL-8, TNF-α, and NGF with statistical difference compared to the non-stressed NPC (nsNPC) control group, but no statistically significant increase in the expression level of CTGF was observed (Fig. 3F).

### 3.3 Stressed NPCs show comparable transcriptome to the patient-derived pain-associated NPC subtype

We compared the single-cell transcriptomes of NPCs before and after the application of combinatory stressors and the NPC1 subtype derived from bpIV0s (nsNPC vs. sNPC vs. bpNPC1). The aNPC data originally used in Fig. 2 was also included as a reference (Fig. S4A). We found that the phenotype of sNPC was comparable to bpNPC1 (Fig. 4). Specifically, both the sNPC and bpNPC1 highly expressed MMP3 yet the nsNPC expressed almost no MMP3 (Fig. 4A). The NPC1 cell population account for 2.7% of nsNPCs, l5.05% of sNPCs, and 8.95% in bpNPCs (Fig. 48). The sNPC cells showed a higher number of commonly shared genes with bpNPC1 (792 genes) than all bpNPCs (496 genes, Fig. 4C). The CP (Z-score) also shows the transcriptomic similarity of sNPC (43.36) to bpNPC1 (68.82), whereas the nsNPC scored the lowest at – 56.2 (Fig. 40). All this evidence suggests higher similarity between sNPC and bpNPC1 than nsNPC.

**Fig. 4.**
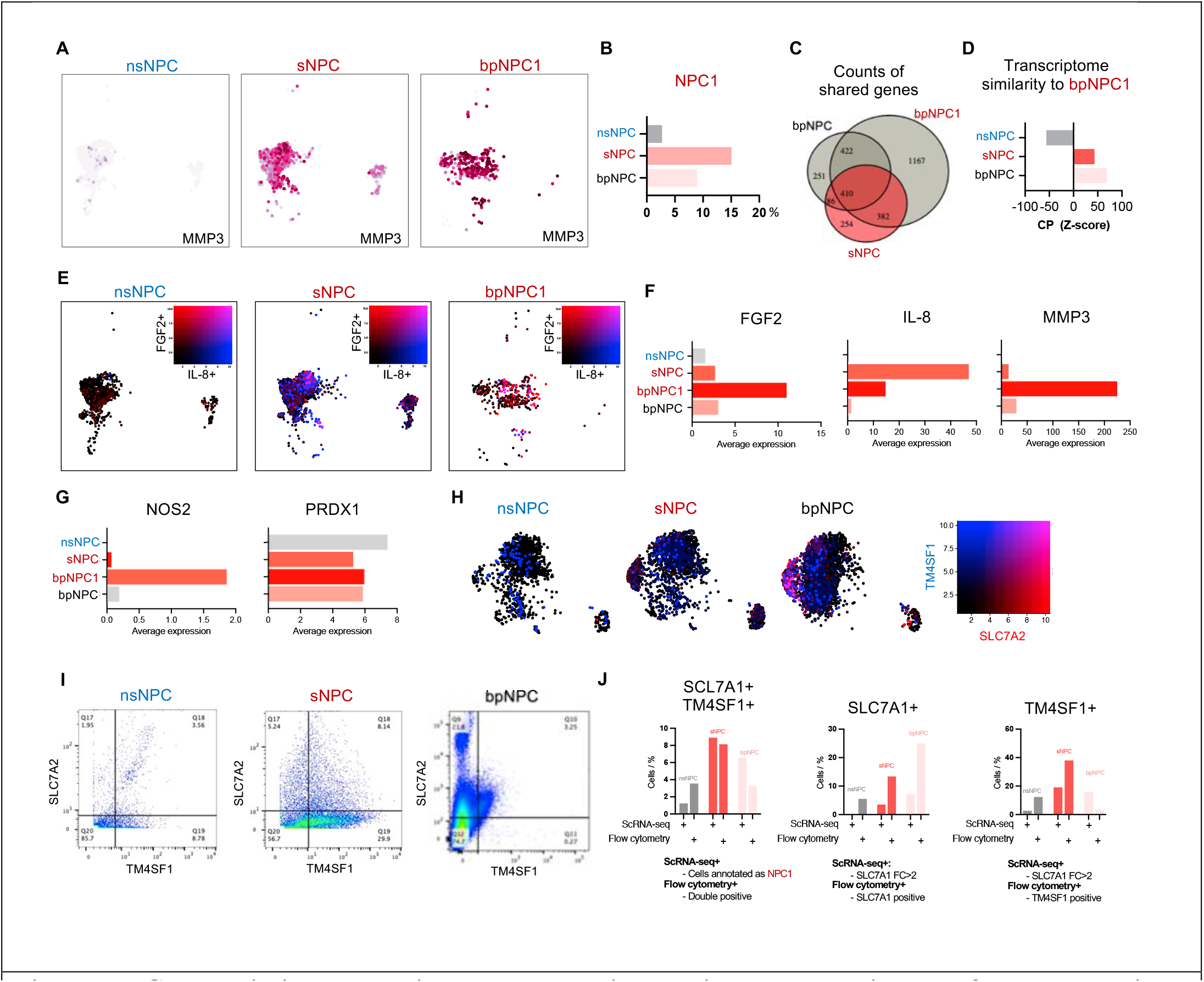
sNPC show similar transcriptome to the pain-associated subtype isolated from LBP patients (bpNPC1). **A)** Gene expression of NPC1 marker, MMP3, is projected on UMAP for single cells in nsNPC, sNPC, and bpNPC1. **B)** Cell percent of NPC1 subtype. **C)** Numbers of shared genes between samples. **D)** Similarity scores of transcriptomes of samples when compared to the bpNPC1. **E)** Co-expression of two secondary NPC1 markers, FGF2 (red) and IL-8 (blue) on UMAP. **F)** Average expression of FGF2, IL-8, and MMP3 markers and **G)** NOS2 and PRDXl markers in nsNPC, sNPC, bpNPC1, and bpNPC. **H)** Co-expression of surface marker TM4SFl (blue) and SLA7A2 (red) are projected on UMAP and used for labeling the pain associated NPC1 cells experimentally. **I)** Flow cytometry of NPC1 using dual surface markers TM4SFl and SLA7A2 in nsNPC, sNPC, and bpNPC. **J)** Cell percentages of the pain-associated NPC1 subtype in the population of nsNPC, sNPC, or bpNPC acquired by either the NPC1 annotation in scRNA-seq or the immunofluorescence-labeling using TM4SFl and SLA7A2 in flow cytometry (marker threshold: fold change >2, expressed both or either gene).

Despite the similarity, we also found differences between sNPC and bpNPC1 in their FGF2 and IL-8 expression (Fig. 4E). IL-8-expressing cells were prevalent in sNPC, whereas FGF2-expressing cells were found in the bpNPC1. Both cell types included a group expressing both genes with much higher expression than nsNPC (Fig. 4E-F).

We also identified highly expressed genes relevant to oxidative stress in the NPC subtypes. We found NOS2 is upregulated in sNPC and bpNPC1 and PRDXl is downregulated in sNPC and bpNPC1 (Fig 4G). Notably, the NOS in sNPC is much lower than bpNPC1. We also found HIF-la, a known marker for hypoxic stress (*59, 60*) highly expressed in sNPC and bpNPC1 compared to nsNPC (Fig. S4B). TRPV4, a known pain marker (*6*) was upregulated in sNPC and bpNPC1 compared to nsNPC (Fig. S4C). The expression of HIF-la and TRPV4 was higher in bpNPC than bpNPC1, sNPC, and nsNPC (Fig. S4B-C). ASIC3, an acidic environment-associated marker (*7*), was expressed higher in bpNPC than aNPC, sNPC, and nsNPC (Fig. S4D).

To validate the hypothesis that there is a common pain-associated cell subtype in both bpNPCs and sNPCs, we selected two surface markers highly expressed in NPC1 cluster, TM4SFl and SLC7A2 (Fig. S3E). Using flow cytometry, we analyzed the presence of double positive TM4SFl+ and SLC7A2+ population in the patient-derived bpNPCs and compared to sNPCs and nsNPCs (Fig. 4I). Flow cytometry confirmed the presence of pain-associated subtype defined by double positive staining in both sNPCs and bpNPCs (Fig. 4J). The cell percent of double positive cells in sNPCs is similar to the cell percent of the pain subtype annotated as NPC1 in scRNA-seq (Fig. 4J). The cell percent of the pain-associated subtype acquired by both methods is higher in sNPCs than bpNPCs and nsNPCs acquired by the same method (Fig. 4J), which indicates that environmental stress can transform NPCs into the pain-associated phenotype (Fig 4J). If only one marker was used in flow cytometry, however, the labeling was inconsistent across the two methods in sNPCs (Fig. 4J). In addition, the SLC7A1 labeled greatly more cells in bpNPCs than TM4SF1 using flow cytometry (Fig. 4J). Double-positive staining should be more accurate than using a single marker.

### 3.4 Stressed NPCs exhibited sensory neuron-inducing function *in vitro*

In a two-step procedure iPSCs were differentiated into neural crest cells and then into iPSC-derived sensory neurons, nociceptor-like cells (iNOCs, Fig. 5A). Successful differentiation was confirmed by immunofluorescence staining for neuronal markers (*6/*) TUBB3 and ISL-1 (Fig. 5B) and RT-qPCR for PRPH, ISL-1, TUBB3, and Brn3a which were statistically higher in iNOCs than iPSCs (Fig. 5C).

**Fig. 5.**
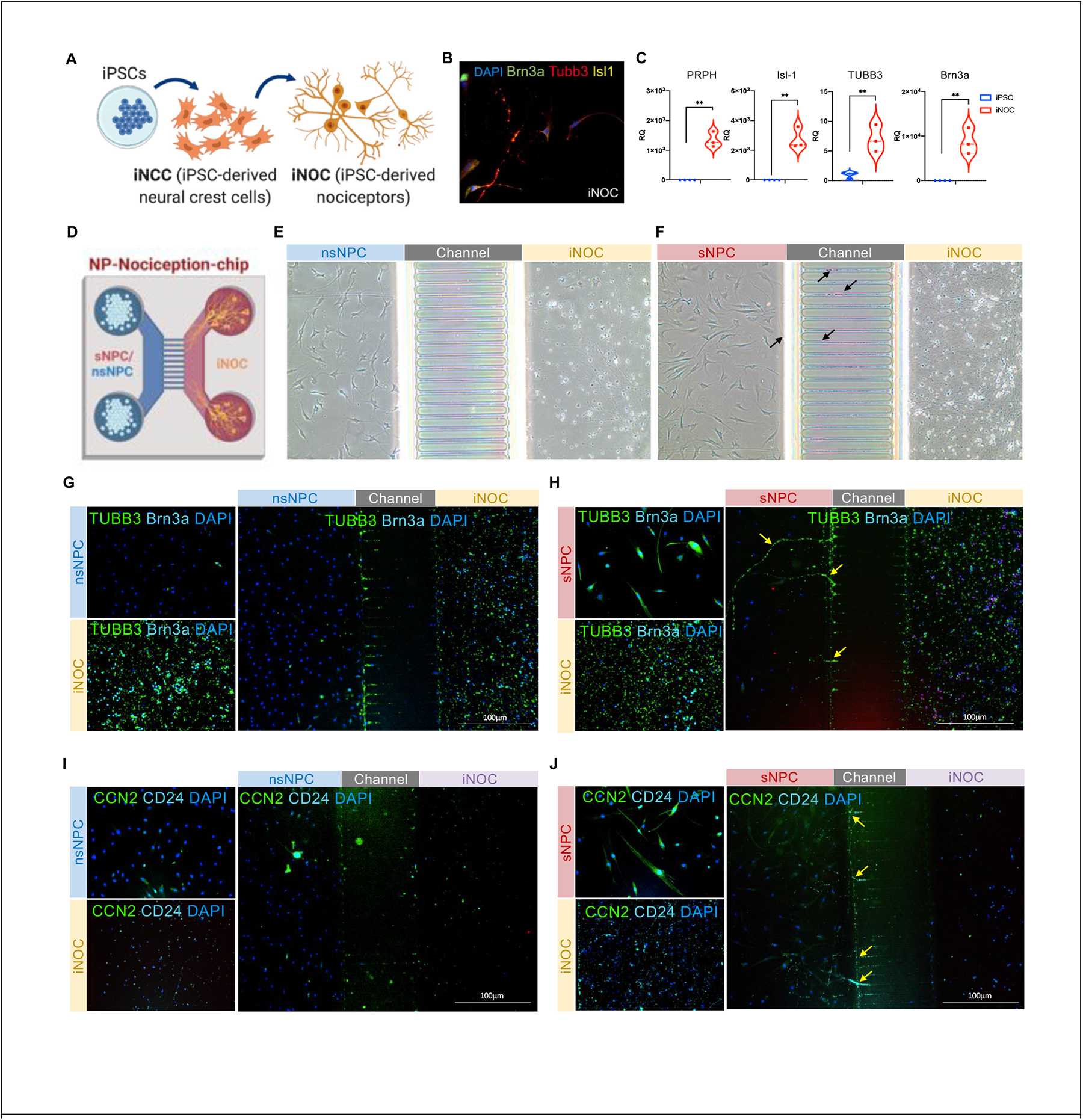
sNPC but not nsNPCs induced nociceptors’ ingrowth in a microfluidic device. **A)** Scheme for iPSC differentiation into induced sensory neurons, nociceptor-like cells (iNOCs) in a two-stage procedure. **B)** Immunofluorescence staining of neuronal markers in iNOC including TUBB3 (green), ISL-1 (white), and DAPI for cell nuclei (blue). **C)** Gene expression levels of neuronal markers PRPH, ISL-1, TUBB3, and Pou4fa (n=4 for iPSC and n=3 for iNOC, \**p*<0.05, \*\**p*<0.01, \*\*\**p*<0.001, \*\*\*\**p*<0.0001). **D)** Illustration showing the co-culture of iNOC and nsNPC or sNPC in a microfluidic chip. Optical images of the co-culture between iNOC and **E)** nsNPC or **F)** sNPC, with black arrows denoting visible axon-like structures traversing the channel. **G-H)** Immunofluorescence images of TUBB3 (green), Brn3a (teal), and DAPI (blue) of iNOC co-cultured with **G)** nsNPC or **H)** sNPC. **1-J)** Immunofluorescence images of CCN2 (green), CD24 (light blue), and DAPI (blue) of iNOC co-cultured with **1)** nsNPC or **J)** sNPC.

The ability of the nsNPCs or sNPCs to induce nerve ingrowth was tested in a co-culture experiment with iNOCs in a microfluidic device that allows crosstalk between the two cell types and crossing over of axons through the microchannels in the middle but prevented cells from crossing over (Fig. 5D). In the nsNPC-iNOC co-culture, we observed no axon ingrowth through the channels to the nsNPC side (Fig. 5E). In the sNPC-iNOC co-culture, we observed several visible axon-like structures crossing the microfluidics channel to the NPC side of the chip (pointed by black arrows, Fig. 5F). We can conclude that the sNPCs effectively induce axon ingrowth from iNOCs compared to the nsNPCs.

We examined the immunofluorescence staining of neuron marker TUBB3 (*62, 63*) and sensory neural marker Brn3a. We found that both markers were almost absent in the nsNPCs but were strongly expressed by the iNOCs (Fig. 5G). We observed the presence of axon-like of TUBB3+ and Brn3a+ signal patterns on the side of the microfluidic chip that included sNPCs and the absence of such structures in the nsNPC cultures, implying the ingrown neural axons traversing the channel in sNPC-iNOC co-culture (Fig. 5G-H). Interestingly, in the sNPC-iNOC co-culture, TUBB3 and Brn3a were strongly expressed in both sNPCs and iNOCs, indicating a crosstalk or sensitization of the NPCs cells by the iNOC cells (Fig. 5H). We also examined the immunofluorescence staining of NPC markers CCN2 and CD24, and both were expressed, as expected (Fig. 5I-J).

### 3.5 Stressed NPCs injected into a healthy rat IVD induced the discogenic pain outcomes *in vivo*

Animal experiments were performed to evaluate whether the sNPC can induce discogenic pain after injection into healthy IVDs in a rat model using a needle size that doesn’t cause IVD degeneration (*47*). Gene expression analysis on day 7 after injection showed that IVDs injected with sNPC highly expressed human MMP3, IL-8, IL-6 with statistical significance when compared to nsNPC group (Fig. 6B), indicating persistence of the stressed phenotype in sNPCs observed *in vitro* (Fig. 3F). Human ACAN expression shows no statistical difference between nsNPC and sNPC groups, suggesting the survival of both injected human sNPCs and nsNPCs (Fig 6B), but was not detected in the saline-injected or narve IVDs. Gene expression analysis using rat primers showed that host IVD cells significantly upregulated expression of inflammatory markers (MMP3, TNF-α, and IL-6) when exposed to sNPCs compared to the nsNPCs, saline control, or narve IVD (Fig. 6C), indicating a paracrine effect in response to the injected sNPCs.

**Fig. 6.**
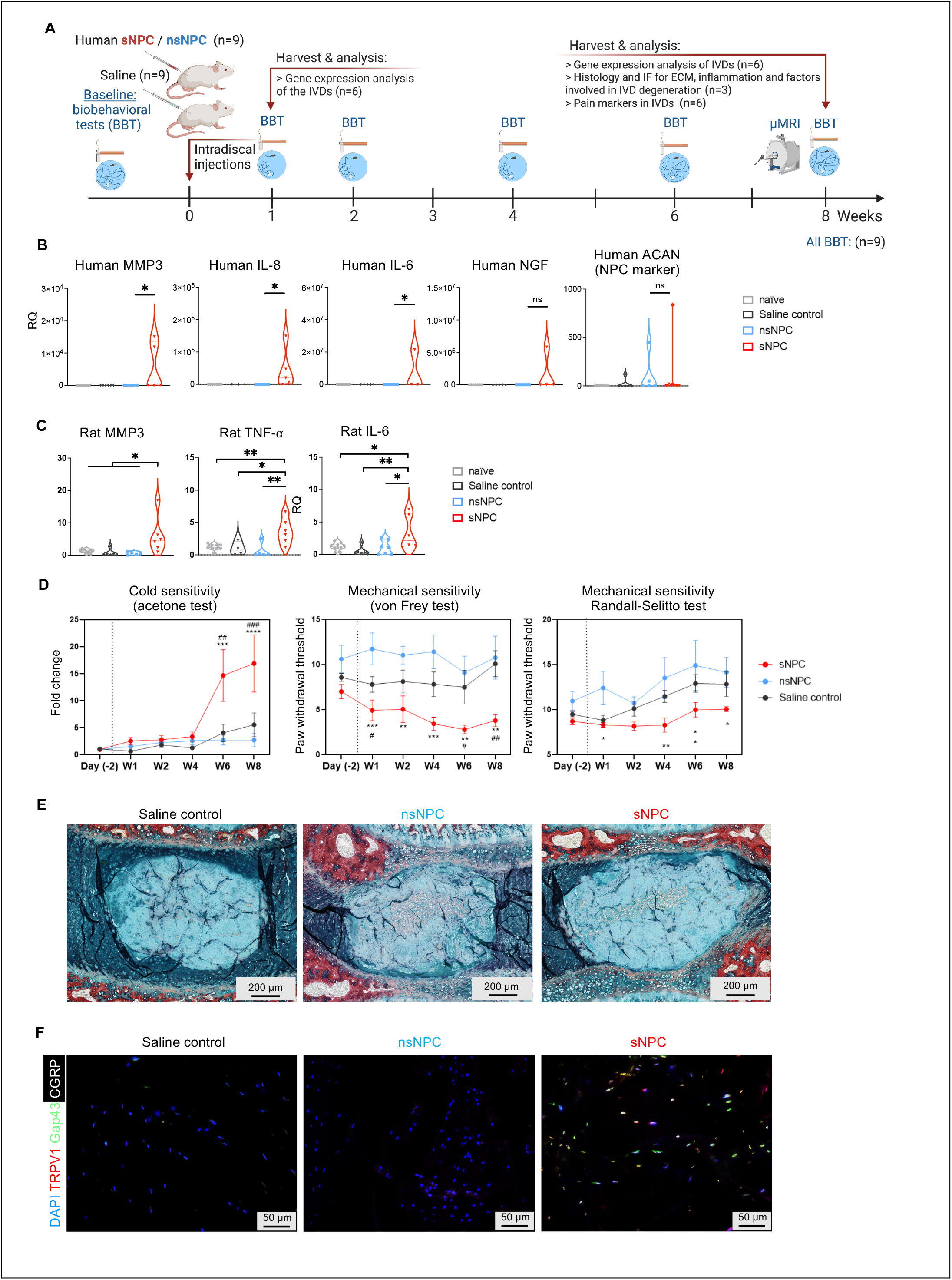
sNPC induced stressful and painful responses after intradiscal injection in rats. **A)** Scheme for group design and timeline for animal experiments in a healthy rat. Gene expression levels of **B)** MMP3, IL-8, IL-6, NGF, and ACAN with human primers and **C)** MMP3, TNF-α, and IL-6 detected with rat primers are shown in RQ (n=6) on day 7 post-injection for rats injected with saline, nsNPCs, or sNPCs. (\**p*<0.05, \*\**p*<0.01, ns or not labeled *p*>0.05). **D)** Biobehavioral test (BBT) pain-related outcomes in Acetone test, von Frey test, and Randall-Selitto tests at 2 days prior to injections, week 1, 2, 4, 6, and 8 post injections. (n=9; \**p*<0.05, \*\**p*<0.01, \*\*\**p*<0.001, \*\*\*\**p*<0.0001 compared to saline control; #*p*<0.05, ##*p*<0.01 compared to nsNPC group). **E)** Histological staining of L4/L5 and L5/L6 levels 8 weeks post injections for saline-injected, nsNPC-injected, and sNPC-injected rat IVDs (Scale bar = 200μm). **F)** Immunofluorescence images of IVDs harvested at week 8 post injections (DAPI (blue), TRPV1 (red), Gap43 (green), and CGRP (white); scale bar = 50 μm).

The biobehavioral tests (BBTs) measuring cold sensitivity (acetone test) and mechanical sensitivity (von Frey and Randal-Sellito tests) were used as indices of an LBP phenotype. Our results showed induction of cold and mechanical hypersensitivity in the sNPC group (Fig. 6D), but not in nsNPC or saline-injected groups. Specifically, the sNPC-injected rats showed statistically increased cold sensitivity in the acetone test at week 6 and 8 post-surgery compared to nsNPC-injected and saline-injected rats. The sNPC-injected rats showed statistically lower paw withdrawal thresholds, indicative of hypersensitivity, in the von Frey test as early as one-week post-surgery compared to saline-injected rats that persisted for 8 weeks post-surgery compared to nsNPC-injected rats. The sNPC-injected rats showed statistically lower paw withdrawal thresholds, consistent with mechanical hypersensitivity, in the Randall-Selitto test at week 1, 4, 6, and 8 post-surgery compared to nsNPC-injected rats. No statistical difference was found between sNPC-injected rats and saline-injected rats at any time point.

Histological analysis of injected IVDs (L4/5, L5/L6 levels) of all tested groups showed no visible disc degeneration compared to reported degeneration models (*47, 64, 65*) 8 weeks post intradiscal injection (Fig. 6E). No visible loss of integrity of the IVDs or matrix degeneration was observed. The µMRI imaging also showed no visible disc degeneration compared to baseline and saline control (Fig. S5). However, the sNPC-injected IVDs showed positive immunostaining for known pain markers: TRPVl, Gap43, and CGRP (Fig. 6F), suggesting that sNPCs can induce upregulation of pain-associated genes 8 weeks after intradiscal injection into a healthy rat IVD. The nsNPC-injected IVDs and saline-injected IVDs did not show positive staining of these pain markers (Fig. 6F), indicating that this upregulation of pain markers was not induced by liquid or human cells injection, but a specific response to the environmentally stressed cells.

## DISCUSSION

This study explored the mechanism of discogenic LBP induction mediated by a subtype of NPCs. The newly described NPC subtype (NPC1) was found to be associated with the occurrence of discogenic LBP in degenerating human IVDs (Fig. 2D). NPC1 was implicated in discogenic LBP induction via upregulated nociceptive and neurogenic functional pathways (Fig. 2H,J), stimulated cell stress, and reactive oxygen species production (Fig. 2H,K). We simulated the NPC1 generation via *in vitro* model of environmental stress application (Fig. 3A,E) and demonstrated that this simulate cell subtype can induce discogenic pain outcomes in an animal model without inducing IVD degeneration.

We recapitulated a degenerative and stressful environment *in vitro* that can stimulate NPCs to show an NPC1-like phenotype expressing inflammatory and degenerative markers (Fig. 3F). The transcriptome of sNPCs was comparable to NPC1 cells derived only from bpIVDs, i.e., bpNPC1 cells, (Fig. 4A-G) and showed that these cells can induce neuronal ingrowth *in vitro* (Fig. 5) and discogenic LBP *in vivo* (Fig. 6). These data suggest that the NPC1 subtype, naturally present in human IVDs (bpNPC1), may be responsible for the onset of early discogenic LBP. This hypothesis is consistent with a study by Richardson et al. which showcased that degenerating human NPCs can induce neurite outgrowth in *vitro* (*35*). Here we extended these results by testing this phenomenon *in vivo* and by addressing the heterogeneity of NPC populations (*35*) as suggested by other studies (*33, 40, 41*).

In this study, we used *in vitro* environmental stressors to model the bpNPC1 cellular phenotype. Direct isolation of the bpNPC1 subtype for these gain-of-function experiments posed several challenges. First, while we were able to label the bpNPC1 subtype with dual surface markers TM4SF1 and SLC7A2 in flow cytometry (Fig. 4H-J), we were unable to establish a reliable method of isolation of this subtype without losing its stress phenotype. Second, *in vitro* culture significantly changed the gene expression profile of bpNPC1 cells, “relaxing” the bpIVD1 cells under regular culture conditions. *In vitro* culture has been reported previously to change cell phenotypes (*66*). Our scRNA-seq of aNPCs after cryopreservation, thawing, and expansion *in vitro* demonstrated a significant change in single-cell atlases (Fig. S4A).

We therefore designed a process (Fig. 3A,E) for recreating the environmental stress that leads to IVD degeneration, aiming to generate a stressed phenotype of NPCs that are functionally comparable to the bpNPC1 subtype. Our optimized combinatory stressors transformed the expanded *in vitro,* non-stressed phenotype of nsNPCs into a stressed phenotype of sNPCs (Fig. 3E-F, 4A-G), which was comparable to bpNPC1 cells (Fig. 4A-G). The effectiveness of this method was shown in several ways. First, the sNPCs and bpNPC1 cells both highly expressed MMP3 (Fig. 3F, 4A,B,F) which is the top marker identifying NPC1 subtype (Fig. 2C and 2D), and more precisely, labeling bpNPC1 cells (Fig. 2F1-G). Second, after applying the stressors *in vitro*, the transcriptome of nsNPCs was transformed towards the bpNPC1 cells (Fig. 4D). Third, the presence of NPC1 (MMP3+) phenotype was confirmed in both sNPCs and bpNPCs in flow cytometry (Fig. 4J). Forth, the sNPCs showed strong expression of inflammatory and degenerative markers (MMP3, IL-1β, IL-8, TNF-*α*) and the neurogenic marker NGF (Fig. 3F), suggesting a strong association with discogenic LBP. Since previous studies harvest the entire NPC populations from degenerative IVDs to study *in vitro* (*35*). This is the first report of the successful generation of a stressed NPC phenotype functionally comparable to bpNPCs or NPC1 (MMP3+) subtype.

In degenerating IVDs, immune cells secrete NGF and BDNF to recruit neural ingrowth and generate pain (*5*). In our study, NPCs demonstrated the ability to play the role of immunomodulating cells. Both naturally present bpNPC1 cells (Fig. 2H) and *in vitro* generated sNPCs (Fig. 3F) highly expressed NGF. Their expression of BDNF was limited (Fig. S3J), but we observed FGF2 overexpression (Fig. 2H,J). The enriched pathway analysis also supports that the bpNPC1 cells could drive neuronal innervation and the expression of both NGF and FGF2 (Fig. 2H), which differs from the previous literature showing NGF and BDNF as co-regulators (*7, 35*). We showed that the sNPCs induced nociception both *in vitro* (Fig. 5) and *in vivo* (Fig. 6). In line with these findings, Richardson et al. reported that degenerating NPCs can induce neurite outgrowth and the presence of NGF is necessary (*35*). FGF2, a growth factor involved in the formation of granulation tissues (*5*), here was expressed in both bpNPC1 and in the sNPC (Fig. 4F). It was also reported that the inhibition of FGF2 can attenuate the mechanical allodynia in a rat model (*67*). Other studies emphasize the catabolic effects of FGF2 which inhibits the synthesis of proteoglycan in a dose-dependent manner (*68*). Similarly, we observed a correlation between FGF2 and MMP3 in our study (Fig. 4E,G). This suggests that FGF2 may play a role in the catabolic process and IVD degradation that warrants further investigation.

Multiple environmental stressors are likely necessary for the onset of NPC-mediated discogenic LBP. When applying environmental stressors *in vitro*, no single stressor was able to stimulate NPCs to express NGF (Fig. 3D). It was the combinatory stressors (IL-lβ, low pH and low glucose) that stimulated strong NGF expression in NPCs (Fig. 3F). The pro-inflammatory cytokine IL-lβ alone induced the expression of inflammatory cytokines in NPCs but did not induce NGF expression, even in high concentration (Fig. 3D). The acidic environment alone induced IL-6 and IL-8 expression in NPCs, but upregulation of NGF was not statistically significant (Fig. 3D). ASIC3, which senses acidic pH, was previously proposed to correlate with pain (*7*). We observed the expression of ASIC3 in bpNPCs was l.8 times higher than in aNPCs, but the overall expression was still very low (Fig. S4D). We also found that the sNPCs generated using combinatory stressors induced LBP phenotype without visible matrix degeneration in rats (Fig. 3F, Fig. 6). Taken together, the environmental stressors in degenerate IVDs may induce the transformation of normal NPCs into stressed, pain-triggering NPCs.

Our findings demonstrate that the NPC-mediated pain cascade following environmental stress may occur without the presence of immune cells. Our proposed NPC-mediated mechanism is compared to the immune cell-mediated hypothesis in Fig. 7. It was previously shown that matrix degeneration stimulates vascular ingrowth and permits the infiltration of immune cells secreting nitric oxide and pro-inflammatory cytokines, including IL-6, IL-8, TNF-*a*, and IL-1B (*5*). Some studies suggest the cytokines secreted by immune cells lead to IVD degeneration (*69*) and discogenic LBP (*70, 71*). We provide evidence that similar inflammation-pain cascade can be initiated by NPCs, circumventing the immune cells. In the current study the bpNPC1 cells expressed inflammatory factors and neurogenic factors (Fig. 2H,J). Their equivalent, sNPCs, were shown to induce neural ingrowth and nociceptive response without the presence of immune cells (Fig. 5). Studies suggest that the nerve fibers can signal pain when inflammatory cytokines are present (*6, 30*). Collectively, the development of discogenic LBP may be independent of immune cells but instead associated with the NPC1 subtype, which has not been previously reported. Previous NPC-focused studies mostly considered the anabolic function of NPCs (expression of proteoglycan and collagen markers such as ACAN and COL2Al) (*31, 72, 73*) rather than pain induction. Also, the precise identification of the pain subtype among NPC population was only made possible in recent years following the advent of scRNA-seq technology (*40, 41, 74*). The NPC-mediated pain mechanism reported in the present study should be further investigated and leveraged for NPC-focused cell therapies.

**Fig. 7.**
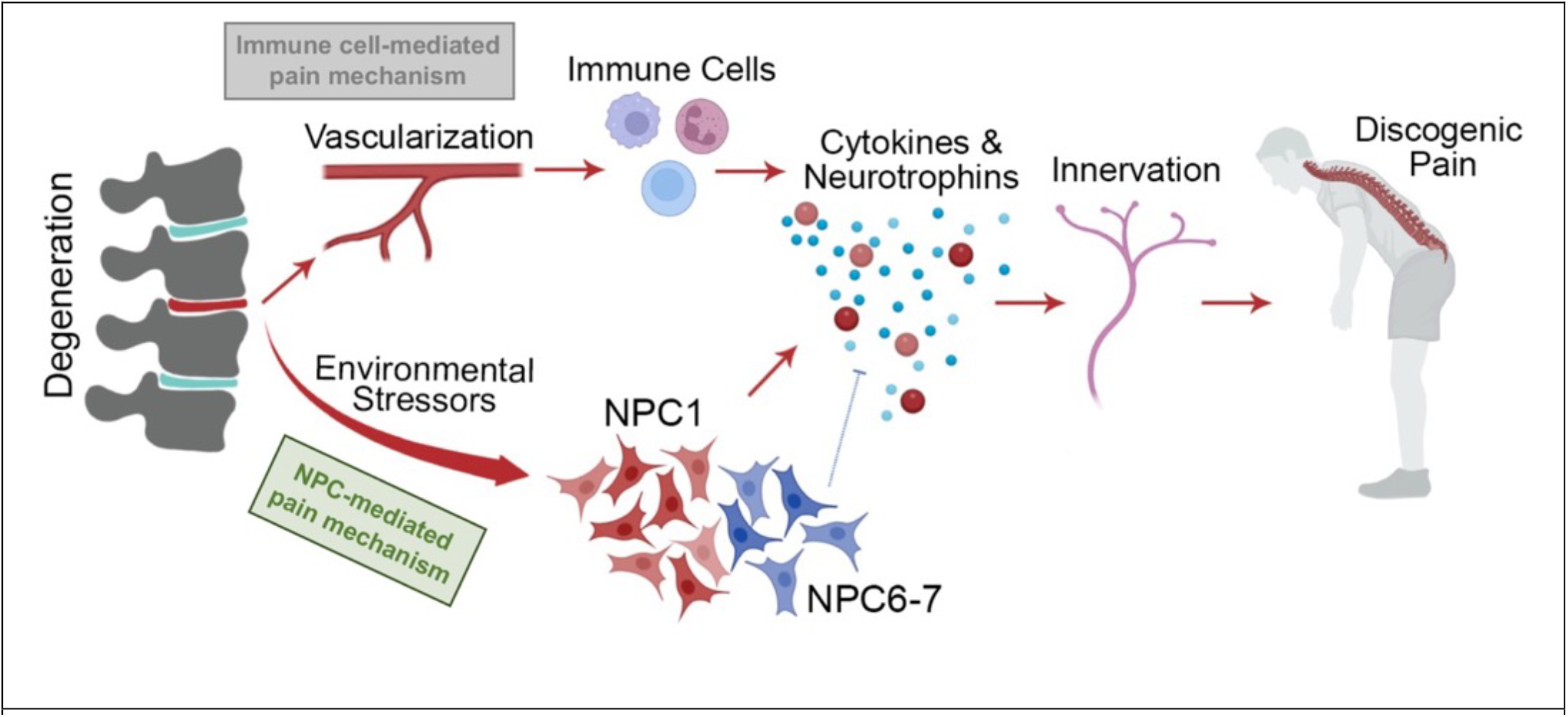
Illustrative summary comparing two pain mechanisms. Immune cell-mediated pain mechanism is an existing theory and NPC-mediated pain mechanism is a newly proposed mechanism in the present study.

We found PRDXl, a gene playing a critical role in reversing cell stress and ROS activity, to decrease with the progression of LBP development (Fig. 2H), whereas an opposite trend along the pseudo-time was found in NGF expression (Fig. 2L). It negatively correlates with bpNPC1 cells but positively correlates with aNPC6 cells (Fig. 2H,J), suggesting that the loss of antioxidant capability in NPCs correlates with the onset and progression of discogenic LBP. These data also suggest that aNPC6 cells, which are enriched in aIVDs, may play a role in mitigating the NPC stress. The top marker for NPC6 subset is PRG4 (lubricin, Fig. 2C,D), which plays role of synovial fluid lubrication (*75, 76*). Together with PRDXl overexpression in NPC6, these findings suggest aNPC6 is a promising cell subtype to relieve the effects of stressful environment on cells.

The aNPC7 subpopulation expressed the nerve growth inhibitor SEMA3A (Fig. S3I) suggesting the potential to inhibit neural ingrowth (*7*). The aNPC7 also highly expressed mitochondrial genes (MTs, Figs. 2C, D, S3D), which is evidence of elevated mitochondrial function. This mitochondrially functional subtype negatively correlates with LBP. NPCs naturally resides in a hypoxia environment (*59, 60*). We also examined the hypoxia-related gene marker, HIF-l*a*, and found it differentially expressed in clusters of bpNPCs or aNPCs (Fig. S3F). In our *in vitro* stress model, the normoxic condition was initially applied at the nsNPC stage, and then the hypoxic condition (2% 0_2_) was applied at the sNPC stage (Fig. S4B). In a hypoxic environment, HIF-1*a* converts oxidative to glycolytic metabolism, thus slowing down mitochondrial activity, maximizing the efficiency of glucose, and the housekeeping function of NPCs (*59*). Madhu et al. suggest that HIF-1*a* induces mitophagy by modulating the mitochondrial translocation of BNIP3 in NPCs (*60*). Risbud et al. also suggest a potential upregulation of VEGF in a hypoxia environment due to its supporting role in NPC survival (not neovascularization) (*59*). We found a correlation between the upregulation of HIF-1*a* (Fig. S3F), BNIP3 (Fig. S3G), VEGF (Fig. S3H), and mitochondrial gene expression in aNPC7 cells (Fig. 2I, S3D). Hence, the mitochondrially active aNPC7 cells may contributing to glycolytic metabolism and survivability for NPCs in hypoxia conditions. In LBP IVDs, however, this aNPC7 subtype diminished (Fig. 2D). Mitochondrial markers are also relevant to neutrophil activity (Fig. 2I). Neutrophils have been shown to induce positive inflammation that protects against the development of chronic LBP (*77*). The aNPC7 subtype may therefore have a protective effect against chronic LBP. Further studies should focus on the potential of NPC6 or NPC7 subtypes in rescuing the pain phenotype of NPCs and attenuating discogenic LBP.

The current study is limited by its scope, thus the potential of the NPC6 and NPC7 subtypes as therapeutic candidates for discogenic LBP has not yet been examined. An additional limitation is the current focused on NPCs; other cells, such as AFCs and their role in LBP induction, should be investigated. In particular, the interplay between AFCs and sensory neurons are of considerable interest to fully decipher the pathology and etiology of discogenic LBP. Finally, the *in vivo* studies were terminated at 8 weeks following IVD injection. While no signs of degeneration were observed at this time point, it is possible that degeneration may be observed at later time points.

Reversing the course of IVD degeneration is challenging and to date no randomized controlled trials have shown superiority over the placebo (*78*). The NPC-mediated pain mechanism proposed in this study suggests an alternative strategy that may attenuate discogenic LBP without the need for regenerating IVD tissues. Specifically, these data provide a paradigm to modulate IVD cells without involving immune cells, by focusing on rescuing NPC phenotype and leveraging the protective subtypes of NPCs (Fig. 7). Precisely targeting the “bad” cell subtype or supplementing the “good” cell subtype in NPC populations may provide more useful strategies for treating discogenic LBP with cell-based therapy.

## MATERIALS AND METHODS

### Study design

The objective of this study was to investigate the role of NPC subtypes in the onset of discogenic LBP. As shown in Fig. 1, the study has been conducted in several different stages: (i) performed the scRNA-seq analyses of human IYO tissues, (ii) Stressing NPCs *in vitro* to recreate the pain subtype and compare it with naturally present pain phenotype, (iii) evaluated the effects of the recreated pain subtype in activating sensory neurons *in vitro*, and (iv) assessed the role of the recreated pain subtype in discogenic LBP in rat models *in vivo*.

(i) The human subject study was approved by the CSMC IRB Pro00020562/ CR00012559. Human IYO tissues were obtained as surgical discards or from cadavers. Informed consent was obtained from the patients undergoing surgery in accordance with the approval of the Institutional Review Board (IRB) at Cedars-Sinai Medical Center (Pro00020562/CR00012559). Specimens were categorized based on patients’ history and diagnoses: non-IYO related spinal symptoms or lack of such as asymptomatic IYO (aIYO, n=4) vs. back pain symptoms primarily associated with degenerated IYO (bpIYO, n=3). For the scRNA-seq analysis, the 7 samples were single-cell sequenced individually and integrated into one Seurat object.

(ii) The IYO cells harvested from aIYO were 2O cultured, and screened with single stressors (pH, glucose, IL-1B, and cyclic mechanical loading) to identify stressful but viability-maintaining conditions. Combinatory stressors were eventually applied to generate the stressed NPCs (sNPCs) One batch of non-stressed or stressed cells was collected for scRNA-seq and the clusters were compared to aIVD and bpIVD samples sequenced in the previous section.

(iii) Stressed or non-stressed NPCs (sNPCs or nsNPCs) were co-cultured with iPSC-derived nociceptors in a microfluidic chip. Details are provided in the supplemental information.

(iv) Appropriate Institutional Animal Care and Use Committee (IACUC) approval (protocol No. 0008089) was obtained before all experiments in this study. Sprague Dawley rats, age 8-10 weeks old, were split into three cohorts: saline, nsNPC, and sNPC (n=9 for each group). Using a 27 Gauge needle, 8µL of saline or cells suspended saline (4 million cells per mL) were injected into the disc at levels L4-L5 and L5-L6. Independent investigators that were blinded to the experimental conditions conducted the biobehavioral tests. Mechanical sensitivity was accessed using von Frey and Randall Selitto biobehavioral testing pre-operatively and post-operatively at time points week 1, week 2, week 4, week 6 and week 8 (n=9). Cold sensitivity was assessed using the acetone test at the same time points. The cohort of rats was randomized, and the experimenter was masked before biobehavioral tests. At sacrifice, the IVD levels L4-L5, L5-L6, and L6-S1 were explanted and collected for RNA extraction and Real-Time qPCR (n=6), for Hematoxylin and Eosin (H&E) staining to evaluate the morphology of the discs (n=3), and for immunofluorescent staining for pain markers (n=6). The experimental methods and materials are detailed in the supplemental information.

## Statistical analysis

For scRNA-seq data, the statistical analyses were performed in R (v4.1.2) using the Seurat package (v4.1.0) or Monocle3 (v1.0.0). For all other data, the statistical analyses were performed in GraphPad Prism (v9.4.1), and *p*<0.05 was used as criteria for significance. All data shown are mean ±SD unless otherwise stated. The two-way analysis of variance (ANOVA) with repeated measures correction was performed. Multiple comparisons test was performed using the Tukey test and the multiplicity-adjusted P value for each comparison was used.

## Supporting information

All Supplementary Materials

## List of Supplementary Materials

### Detailed Materials and Methods

List of supplementary figures:

Fig. Sl. The number of genes detected per cell (nFeature_RNA), reads per gene (nCount_RNA), and the percent of mitochondrial genes (percent.mt) of single-cell data for each sample.

Fig. S2. CellScale^TM^ bioreactor. Optical photos of (A) the components for the CellScale^TM^ bioreactor before assembly and (B) after assembled and placed in an incubator. The samples were placed in the sample holders which were placed in the wells on the base. (C) the compression strain and sensed force applied onto hydrogel construct loaded with NPCs. The frequency is lHz, and the strain is l2.5% and 30% respectively.

Fig. S3. Supplemental figures for single-cell analyses of human bpIVD and aIVD cells.

Fig. S4. Supplemental figures for single-cell analyses of human nsNPC, sNPC, bpNPC, and aNPC cells. Fig. S5. The µMRI images showing rat IVDs injected with sNPCs or saline.

## Acknowledgments

We thank Cedars-Sinai genomics core for the help in scRNA-seq. We thank the Cedars-Sinai Medical Center Research Imaging Core for the support and assistance in this work. We thank Cedars-Sinai Biobank and Research Pathology Resource for providing human samples. We thank the Cedars-Sinai Biobehavioral Research Core.

## Funding

Cedars Sinai Board of Governors Regenerative Medicine Institute

National Institutes of Health grant K01AR071512 (OS)

National Institutes of Health grant R34NS126032 (OS, LSS)

California Institute for Regenerative Medicine for EOUC4-12751 grant (WJ)

## Author contributions

Conceptualization: WJ, JOG, WT, HWB, LSS, OS

Methodology: WJ, JOG, GK, JS, JTW, SS, KS, JLC, WT, PA, CJ, CC, LEAK, TT, JB, OS, TGP, MK, HWB, LSS and OS

Investigation: WJ, JOG, GK, JS, JTW, SS, KS, JLC, WT, PA, CJ, CC, LEAK, TT, JB, OS, TGP, MK, HWB, LSS and OS

Visualization: WJ, JOG, GK, JS, JTW, KS, and OS

Funding acquisition: LSS, OS

Project administration: WJ, JOG, WT, OS

Supervision: JOG, WT, JB, HWB, LSS, OS

Writing – original draft: WJ, JOG, GK, JTW, LSS, OS

Writing – review & editing: WJ, JOG, LEAK, TT

## Competing interests

Authors declare that they have no competing interests

## Data and materials availability

The raw sequencing data in the scRNA-seq are accessible in Gene Expression Omnibus: GSE222182. The original code is accessible on GitHub: https://github.com/jason199112345/scRNA-seq-for-back-pain-study.git. The rest data are available in the main text or the supplementary materials.

## Notes

### Competing Interest Statement

The authors have declared no competing interest.

