## Supplementary material for "Environmentally stressed human nucleus pulposus cells trigger the onset of discogenic low back pain": All Supplementary Materials

### Detailed Materials and Methods

#### Human tissue sample preparation and single-cell RNA sequencing

Discarded human IVD specimens were obtained from 7 donors (Table S1). Informed consent was obtained from the patients undergoing surgery in accordance with the approval of the Institutional Review Board (IRB) at Cedars-Sinai Medical Center (Pro00020562/CR00012559).

**Table S1.** Patient characteristics, specimen descriptions and scRNA-seq process results

| Subject # | 1 | 2 | 3 | 4 | 5 | 6 | 7 |
|---|---|---|---|---|---|---|---|
| Group assignment | aIVD | aIVD | aIVD | aIVD | bpIVD | bpIVD | bpIVD |
| Sex | M | M | M | F | M | M | F |
| Age | 45 | 57 | 45 | 66 | 59 | 70 | 40 |
| IVD levels | L2-L3 | L1-L4 | L5-S1 | L2-L3 | L4-L5 | L4-L5 | C6-C7 |
| IVD-associated pain? | Not reported | Not reported | Not reported | Not reported | Reported | Reported | Reported |
| Source | Cadaver | Cadaver | Surgical Discards | Surgical Discards | Surgical Discards | Surgical Discards | Surgical Discards |
| Cells sequenced | 2722 | 5990 | 508 | 5523 | 1458 | 417 | 3979 |
| Mean reads/cell | 73328 | 97823 | 386409 | 114588 | 136193 | 494169 | 53924 |
| Median genes/cell | 3231 | 3592 | 2427 | 3618 | 1649 | 3288 | 2732 |
| Median UMI/cells | 13124 | 13648 | 9405 | 14672 | 5476 | 14089 | 9156 |

**Isolation of cells from human intervertebral discs**

Human NP tissue specimens were processed, and cells were isolated from tissues as previously reported (*79-81*). Briefly, the isolated tissues were manually minced into ~1mm pieces and washed two to four times with phosphate-buffered saline (PBS). Digestion media (DMEM/F12+1% Antibiotic/Antimycotic) with 2mg/ml of Pronase Protease (Sigma Aldrich, St. Louis, MO, USA, Cat # 53702) was added to the NP tissue and allowed to digest for one hour at 37°C. The tissue was then centrifuged for 10min at 1200 prm for 10 min. The resulting supernatant was discarded, and the remaining tissue was then resuspended in digestion media with 110mg/ml of Collagenase Type-1S (Sigma Aldrich, Cat # C1639), and allowed to digest overnight at 37°C. The resulting liquid was filtered through a 70µm filter and centrifuged for 10min at 1200rpm at room temperature. The supernatant was discarded, and the remaining cells were counted and plated in a 100mm dish at a concentration of ~1 million cells per plate.

**Single-cell RNA sequencing and analyses for nucleus pulposus cells**

**a) Samples preparation for single-cell RNA sequencing**

Four aIVD and three bpIVD samples were processed for single-cell RNA sequencing. Following the digestion and filtration described in section 3.3, the collected cell pellet was resuspended in PBS at a concentration of 1,500 cells/µL and used for single-cell RNA sequencing using the Chromium Single-cell 3' v3.1 Reagent Kits from 10x Genomics and sequenced as reported (*81*). Briefly, we followed 10x Genomics protocols with a target capture of ~3,000 cells captured on a Chromium Controller, and then the libraries were sequenced (28bp Read 1 and 91bp Read 2) with a single sample index (8bp) on an Illumina NovaSeq 6000. The depth is >50,000 raw reads per cell. Bcl2fastq v2.20 (Illumina, San Diego, CA) was used in converting raw sequencing data into fastq format. Cell Ranger 6.0.0 and Loupe Browser 3.0.0 were used for analyzing and visualizing raw sequencing data. The configurations for scRNA-seq are summarized in Table S1. Quality control was performed, and only single-cell libraries that passed quality control filters were aggregated across all experimental batches and analyzed together. Uniform Manifold Approximation and Projection (UMAP) was used as the method for non-linear dimensional reduction of the data. Cells were classified into unsupervised clusters and assigned to different cell subtypes based on known marker expression.

**b) Preliminary data processing, quality control, and dimensional reduction**

The matrix, barcoding, and feature dataset of 7 samples were loaded individually using the Read10X function, then converted into 7 individual Seurat objects (Seurat package v4.1.0). The number of detected genes (nFeature_RNA) and the percent of mitochondrial genes (percent.mt) of each Seurat object are shown in Fig. S1. Quality control has been implemented following these criteria: nFeature_RNA should be greater than 200 and smaller than 7000, and the percent.mt should be smaller than 10%. Cells that passed these criteria were selected for the following analysis. The criteria considered the fact that most cells of each Seurat object fall into the range of 200-7000 in terms of nFeature_RNA and <10% in terms of percent.mt as shown in Fig. S1. After quality control, the 7 Seurat objects were then combined into a matrix file. Next, the normalization was performed for the combined Seurat object. Total 6000 most variable genes were selected for analysis. The dataset was then, anchored, integrated, scaled and centered to remove the batch effects. Principal component analysis was carried out at PC = 30. The dataset was then processed with dimensional reduction using the UMAP method. The Seurat package classified cells into 12 unsupervised clusters at 0.5 resolution. Each cell kept its identity of the sample type (bpIVD or aIVD), cluster number (0-12), and original sample (bpIVD samples 1-3, aIVD samples 1-4), which will be referred to in the downstream analysis.

**c) Cell compositions, markers, gene ontology enrichment, and pathway analyses for nucleus pulposus cells populations using single-cell data**

Clusters with NP cell markers were (NPC1-7) subsetted for further analyses of the combined Seurat object containing all samples. The percent of cells derived from aIVD samples or bpIVD samples were calculated in R based on the identities of sample type and cluster number for each cell. The genes differentially expressed (upregulated or downregulated) in one cluster than other clusters were identified for each cluster. Among those differentially expressed genes, the one with the highest fold change (positive) and $p > 0.0001$ was selected as the marker for this cluster (subtype). The differentially expressed genes (upregulated or downregulated) from aIVD or bpIVD samples were also identified separately. All differentially expressed genes with $p > 0.0001$ for cells from NPC1 clusters and derived from bpIVD (bpNPC1), cells from NPC6 cluster and derived from aIVD (aNPC6), and cells from NPC7 clusters derived from aIVD (aNPC7) were selected for gene enrichment and pathway analysis. The average expression levels of genes of interest were calculated in R. The average expression levels of enriched canonical pathways and biological functions were calculated in Qiagen Ingenuity Pathway Analysis (IPA, version 81348237). The percent of cells derived from bpIVDs or aIVDs has been normalized to the total cell counts calculated in R. In volcano plots comparing bpNPC1 and aNPC1, the threshold of fold change is 2 and $p$ value is $10^{-36}$: markers with fold change > 2 or < -2 ($Log_2$Fold Change > 1 or < -1) and $p$ value < $10^{-36}$ ($-Log_{10}p > 36$) are colored with red and listed in the text box. Other markers were colored gray and not listed. The threshold for bpNPC6 vs. aNPC6 and bpNPC7 vs. aNPC7, the threshold for fold change is 2 and $p$ value is $10^{-3}$.

**d) Pseudo-time trajectory for nucleus pulposus cells using single-cell data**

The pseudo-time developmental trajectory analysis was performed using the Monocle3 (v1.0.0) package. The combined Seurat object was firstly subset to a new Seurat object containing NPC1, 2, 5, 6, and 7 and was then loaded into a Monocle3 object. The Monocle3 package constructed a pseudo-time trajectory and visualized with each subtype assigned to a different color. The expression level of genes of interest along the trajectory or pseudo-time was further visualized.

**e) Single-cell RNA sequencing for stressed and non-stressed nucleus pulposus cells**

The IVD cells harvested from asymptomatic IVD following the same procedure in 3.2 were cultured in 2D culture as non-stressed cells. The non-stressed cells were applied with the combinatory stressor following the procedure in 3.5 which thus were considered stressed cells. One batch of non-stressed and one batch of stressed were sequenced following the procedure in 3.4.1. The same procedure of preliminary data processing, quality control, and dimensional reduction as in 3.4.2 was applied to the non-stressed or stressed cells which were then integrated together with all samples in the combined Seurat object in 3.4.2. The newly combined Seurat object contained cells from 9 samples: 4 aIVDs, 3 bpIVDs, 1 sample of non-stressed cells from aIVD, 1 sample of stressed cells from aIVD. The newly combined object was also subset to NPC subtypes which resulted in a comparison of NPC across different samples: nsNPC from non-stressed samples, sNPC from stressed samples, bpNPC from bpIVD samples, aNPC from aIVD samples. We specifically subset NPC1 (MMP3+) subtype derived from bpIVD samples to form a cell subset of interest: bpNPC1 (MMP3+). The differentially expressed genes, marker, and gene enrichment and pathway analyses of each cluster were identified with the Seurat package (v4.1.0) in R (v4.1.2, same procedure as in 3.4.3). The cell atlas and gene expression levels of genes of interest were calculated and visualized with the Seurat package (v4.1.0) in R (v4.1.2). The comparisons of shared genes and transcriptome similarity were performed in Qiagen Ingenuity Pathway Analysis (IPA, version 81348237).

**Application of exogeneous stressors on nucleus pulposus cells**

NPCs harvested from aIVDs were expanded in 2D culture using normal NPC culture media in standard cell culture conditions (95% air, 5% $CO_2$, 37°C, sterile, humidified). The normal NPC culture media was prepared using Dulbecco's Modified Eagle Medium F12 (1:1) base media (Gibco™ 11320033) added with 10% FBS, 1% antibiotic-antimycotic (Gibco™ 15240062), 50mg/L L-ascorbic acid (Sigma-Aldrich A4544) and 0.05 mM L-glutamine (Gibco™ 25030081). NPCs were first cultured in 2D tissue-treated culture plate till confluency prior to their placement into a hypoxic chamber (2% $O_2$, 5% $CO_2$) and the addition of stressor or non-stressor media. The 2D cultured NPCs (P3) were then subjected to single stressors of either inflammatory cytokines (IL-1β), low pH environment, low glucose, or mechanical compression.

To apply IL-1β stressor, the culture media was modified from normal NPC culture media by adding IL-1β at 0, 0.25, 1.0, 2.5, and 5.0ng/ml. NPCs were cultured with IL-1β added stress media or unmodified media in 2D tissue culture-treated plates for 1 day and tested for viability and gene expression. To apply the glucose stressor, we mixed a base media with high glucose content (Gibco™ 12634010) and a glucose-depleted media (Gibco™ A2494301) to produce a media that had a concentration of 0.1, 0.5, 1.0, and 3.1g/ml glucose. The mixed media was added with 5% FBS, 1% antibiotic-antimycotic (Gibco™ 15240062), and 4mM L-glutamine (Gibco™ 25030081). NPCs were cultured with glucose-modified (glucose = 1, 0.5, 0.1g/ml) or unmodified (glucose = 3.1g/ml) media in 2D tissue culture-treated plates for 14 days for viability and 4 days for gene expression. To apply the pH stressor, we modified the previous mixed media with a 1.0 glucose concentration. We modified the media pH to 6.75, 7.0, and 7.25 using 1N HCl or 1N NaOH, or no modification, and then filtered the media with a 200nm filter for sterilization. NPCs were cultured with the modified or unmodified media (pH = 7.80-7.84) in 2D tissue-treated culture plates and tested for viability and gene expression. To apply mechanical stress, a previously published hydrogel 3D construct (*51*) was used to encapsulate the NPCs. The NPC-loaded hydrogels were placed pre-cultured in 2D tissue culture-treated plates under standard cell culture conditions (95% air, 5% $CO_2$, 37 °C, sterile, humidified) for 1 day, and then loaded into a bioreactor (Fig. S2A-B, CellScale MechanoCulture TX) inside the hypoxia chamber (2% $O_2$, 5% $CO_2$). The pincher will apply deformation to the hydrogel construct from the top as programmed. The bioreactor permitted accurately controlled displacements and recorded the locations and forces on the pincher (Fig. S2C). We programmed the motion of the pitcher for 48h. For every 24h, the first 12h was set to be relaxed status without deformation, from 12-13h, there was 1h dynamic pressing time during which the pincher was pressing down to a deformation of 0%, 6.7, or 12.25% of the total height of the samples at 1Hz frequency (Fig. S2C).

The rest 13h-24h was a relaxed status again without any deformation or movement of the pincher. In total, each sample was kept culture in the bioreactor, under hypoxia condition (2% $O_2$, 5% $CO_2$), without mechanical stress for 1d prior to testing the viability and gene expression.

The *in vitro* viability was tested using CellTiter-Glo® 3D Cell Viability Assay kit for all samples. A plate reader (SpectraMax M3, Molecular Devices) was used to obtain the luminescence reading. The viability was calculated as the percent of luminescence for each sample compared to the luminescence of the control sample, that is 0ng/ml for IL-1β stressor, 7.84 for pH, 3.1g/ml for glucose, and 0% for mechanical loading.

The viability of each sample after single stressor experiments was tested. The viability of each sample after experiments were tested. The gene expression analyses were performed for each sample using RT-qPCR assays. The RNeasy® Lipid Tissue Mini Kit (Qiagen, 1023539) was used for 3D NPC-loaded hydrogel samples, and the RNeasy® Plus Micro Kit was used for 2D cultured NPCs in the pH, glucose, and IL-1β single stressor experiment. The resulting RNA was transcribed into cDNA using a high-capacity reverse transcription kit (Applied Biosystems, MA, USA, catalog #4368813). Gene expression was evaluated using Taqman gene expression assays (Applied Biosystems, MA, USA). All assays were normalized to the 18S housekeeper gene.

The combinatory stressor experiment was performed under hypoxia conditions (2% $O_2$, 5% $CO_2$) in a hypoxia chamber. The NPCs were first cultured in 2D tissue culture-treated plates till confluency with normal NPC culture media. The media was prepared based on the previous glucose-modified media (glucose = 0.1g/L) which was adjusted for pH using 1N NaOH or 1N HCl. After the pH was adjusted to 6.80-6.90, a 200nm filter was used for sterilization. The NPCs were then cultured with the modified media for 3 days. The IL-1β cytokines were then added to the media at 2.5ng/ml. 24h later, collected cells for viability and gene expression using the same method described for the single stressor experiment. The NPCs had been subjected to glucose and pH stressors for a total of 4 days, and IL-1β for a total of 1 day.

**Co-culture of stressed nucleus pulposus cells with iPSC-derived nociceptors in an NP-nociception microfluidic chip**

The iPSCs (83i-ctr22n1) obtained from CSMC iPSC core were differentiated into immature Nociceptors (iNOCs) using the Senso DM kit (Anatomic Inc., Minneapolis, MN) per manufacturer protocol. The microfluidics chips (Xona Microfluidics, Research Triangle Park, NC) were prepared one day prior to iNOC seeding per manufacturer protocol. Chips were additionally coated with a layer of iMatrix511 silk and allowed to incubate overnight (1:100, Nippi Corp., Tokyo, Japan). Immature iNOC were seeded on one side of the chip at a density of ~100,000 cells per two wells. iNOC were fed daily with Senso-MM media (Anatomic Inc.) and allowed to mature. At 7 days post iNOC seeding, the sNPC were seeded on the other side of the chip at a density of 24,000 cells per two wells. The sNPC were fed every two days with stress media. At day 21 post NPC seeding, chips were fixed for 15 minutes with 4% Paraformaldehyde. For immunocytochemistry staining of the chips, non-specific binding was blocked by the addition of normal donkey serum (Jackson ImmunoResearch, West Grove, PA). The chips were stained with primary antibodies and allowed to incubate overnight at 4°C. On day two, the chips were washed 0.1% Tween in PBS and then incubated in the dark for one hour at room temperature with the secondary antibodies. Finally, the chip was washed four times with 0.1% PBS-Tween and then stained with 4',6-Diamidino-2-Phenylindole dihydrochloride (DAPI, 300nM, Invitrogen) for 5 minutes in the dark. Chips were imaged with Revolve (RVL100B2, Echo Laboratories, San Diego, CA).

**Intradiscal injection of stressed nucleus pulposus cells into healthy rat intervertebral discs**

**a) Intradiscal cell injections**

All animal study procedures were approved by the Institutional Animal Care and Use Committee (IACUC) of Cedars-Sinai Medical Center (IACUC0008089). 27 healthy, female Sprague Dawley rats (Charles

River, Massachusetts), (8-10-week-old, 220–240g) were kept in a controlled environment (22 ± 1°C, 50 ± 1% relative humidity, and 12/12h light/dark cycle) for the duration of the study.

Rats were anesthetized (Isoflurane 2-2.5% maintenance), and ophthalmic ointment was placed in the animal's eyes to prevent corneal drying. Before the start of the surgical procedures, buprenorphine (0.1mg/kg) was injected subcutaneously. Hair on the surgical site was clipped using electric clippers. The surgical site was aseptically prepped by thoroughly disinfecting with betadine followed by alcohol in alternating wipes. An abdominal straight incision (~7cm) was made with sterile surgical scissors. The abdominal incision was extended through the linea alba into the abdominal cavity. Note that incisions with surgical scissors, as opposed to a scalpel, reduces bleeding and the risk of damage to the underlying tissues. The intestines were deflected to the rat's right to expose the abdominal aorta and the left kidney. Anatomical landmarks were then palpated to determine the spinal region to be exposed in the upper caudal vertebrae.

Rats were randomly assigned into three cohorts: saline only control, nsNPC and sNPC. Prior to needle insertion, a mini-C-arm was used to clearly identify the L4-L5 and L5-L6 levels. A Hamilton Syringe with a 27G needle was used to deliver 8μL of saline or cells (nsNPC or sNPC) solution in saline (~4k cells per μL) injected into the intervertebral disc. The sNPC cells were prepared following the combinatory stressing protocol mentioned above (pH at 6.75, glucose 0.1g/L, and IL-1β 2.5ng/ml). The nsNPC cells were prepared with the same condition as sNPC cells except for the media was unmodified (pH=7.80-7.84, glucose 3.1g/L, and no addition of IL-1β). After intradiscal injections, the peritoneal contents were replaced, and the rectus fascia, and skin were closed in layers.

Postoperative, warm, approximately 37°C, normal saline was administered subcutaneously and thermal support (heat lamp placed approximately 18 inches from the animal) continued to be provided throughout recovery to protect against hypothermia.

Animals were placed in a new home cage (on a paper towel to prevent bedding from adhering to the ophthalmic ointment and eyes) with fresh bedding and gel diet or water-soaked food on the cage floor on a Petri dish and monitored to ensure adequate recovery from anesthesia. Animals will be observed, and buprenorphine (0.03mg/kg) administered subcutaneously early in the morning on the day after surgery.

Post-operatively, rats were singly housed and received enrofloxacin (SC, 1mg/kg) for up to 3 days. Non-absorbable sutures were removed under physical restraint or brief isoflurane anesthesia 7-10 days after surgery. Once sutures were removed, animals were paired housing for 8 weeks prior to Euthanizations.

**b) Biobehavioral tests of rats following the intradiscal injection of stressed nucleus pulposus cells**

Mechanical sensitivity was assessed using Von Frey and Randall Selitto biobehavioral testing. Rats underwent biobehavioral testing pre-operatively and post-operatively at time points week 1, 2, 4, 6 and 8.

For Von Frey testing, rats were placed onto a gridded platform and the glabrous areas (between the interdigital pads) of their paws were stimulated with Von Frey Filaments (IITC Life Science, Woodland Hills, CA). When the animal is alert, and with all four paws caring the weight in contact with a grid, von Frey filaments (0.8g, 1.3g, 1.8g, 2.4g, 2.9g, 7.5g, 11g, 12g, 13g) were applied to the hind paws in the ascending order of force with the starting point of 2.9g. They were pressed to the specific midpoints of the plantar surface of the hindpaw until the filament just bent. The withdrawal threshold was assessed as the force that produces a positive response. A response was considered positive when the animal exhibits any nocifensive behaviors such as paw withdrawal, licking, shaking, etc., either during the application or immediately after in 3 of 5 consecutive applications within one trial.

The "up-down" Von Frey method (*82*) is used to determine the mechanical force required to elicit a paw withdrawal response in 50% of animals, based on the statistical formula used to determine LD50s (*83, 84*).

For the Randall Selitto test, the rats hind paws were placed onto the Randall Selitto apparatus (Ugo Basile, Gemonio VA, Italy). Increasing mechanical pressure at a constant rate of 16 grams per second from 0 to 250g was applied to the dorsal surface of the hind left and right paw until withdrawal or vocalization occurred. The paw was placed on a small plinth under a cone-shaped pusher with a rounded tip. Each testing session consisted of 3 trials with a 15 min break between the trials. The withdrawal threshold was recorded in grams and the resulting numbers were averaged.

Cold sensitivity was assessed at the same time point using the acetone biobehavioral test. For acetone, rats were placed onto a gridded platform, and ~ 0.1mL of acetone was unilaterally placed onto the plantar surface of the hind left and right paw of rats with 1min interval between the limbs. The rats were observed and videotaped for one minute. The video recordings were played back in slow motion to accurately analyze the response to acetone. To assess cold hypersensitivity, all nocifensive responses were identified and categorized into groups: lifting, flicking, stamping, paw licking, paw sniffing, paw itching, paw holding, ankle motions such as dorsiflexion, paw rotations in and out, and withdrawal reactions. Sensitivity was recorded by quantifying the duration of all nocifensive responses. The duration of all responses was measured for one minute with a digital stopwatch. The total amount of time spent doing the selected behaviors was summed to determine the total amount of time reacting to the stimuli.

**c) Gene expression analysis post IVD injections**

At sacrifice, the IVD levels L4-L5, L5-L6, and L6-S1 and dorsal root ganglia (DRG) levels L1, L2, and L3 were explanted. The harvested tissue was macerated, and total RNA was collected using a RNeasy Lipid Tissue Mini Kit (Qiagen, catalog #74804) following the manufacturer's protocol. The resulting RNA was transcribed into cDNA using a high-capacity reverse transcription kit (Applied Biosystems, MA, USA, catalog #4368813). Gene expression was evaluated using Taqman gene expression assays (Applied Biosystems, MA, USA). All assays were normalized to 18S housekeeper gene.

d) Histology and immunofluorescence

At sacrifice, the whole spines were explanted. The spines were fixed in 4% formaldehyde solution, decalcified in 0.5M EDTA pH 7.4 for four to six weeks, passed through a graded series of ethanol solutions, and embedded in paraffin. Five-micron-thick sections were cut from the paraffin blocks. Hematoxylin and eosin (H&E) staining was performed to evaluate the morphology of the discs. For immunofluorescent staining, tissues were deparaffinized, and the antigens were retrieved by incubation in Proteinase K (Agilent, Carpinteria, CA, catalog #S3020 for 20min at room temperature. Nonspecific antigens were blocked by applying Normal Donkey Serum (Jackson ImmunoResearch, catalog #077-000-121). Slides were stained with primary antibodies. The primary antibodies were applied to the slides, after which the slides were incubated at 4°C overnight and washed using PBS; the slides were then incubated with secondary antibodies for one hour at room temperature. Finally, the slides were stained with DAPI (Invitrogen, catalog #D1306) for five minutes in the dark. ProLong™ Gold Antifade mounting medium (Invitrogen, catalog #P36935) was applied to the tissue. Images were captured using a Carl Zeiss Axio Imager Z1 fluorescent microscope (Carl Zeiss, Oberkochen, Germany) equipped with ApoTome and AxioCam HRc cameras. Negative controls were processed using identical protocols while omitting the primary antibody to exclude nonspecific staining. Images were captured with 4x4 tile scans at 20x objective.

**Supplementary Figures**

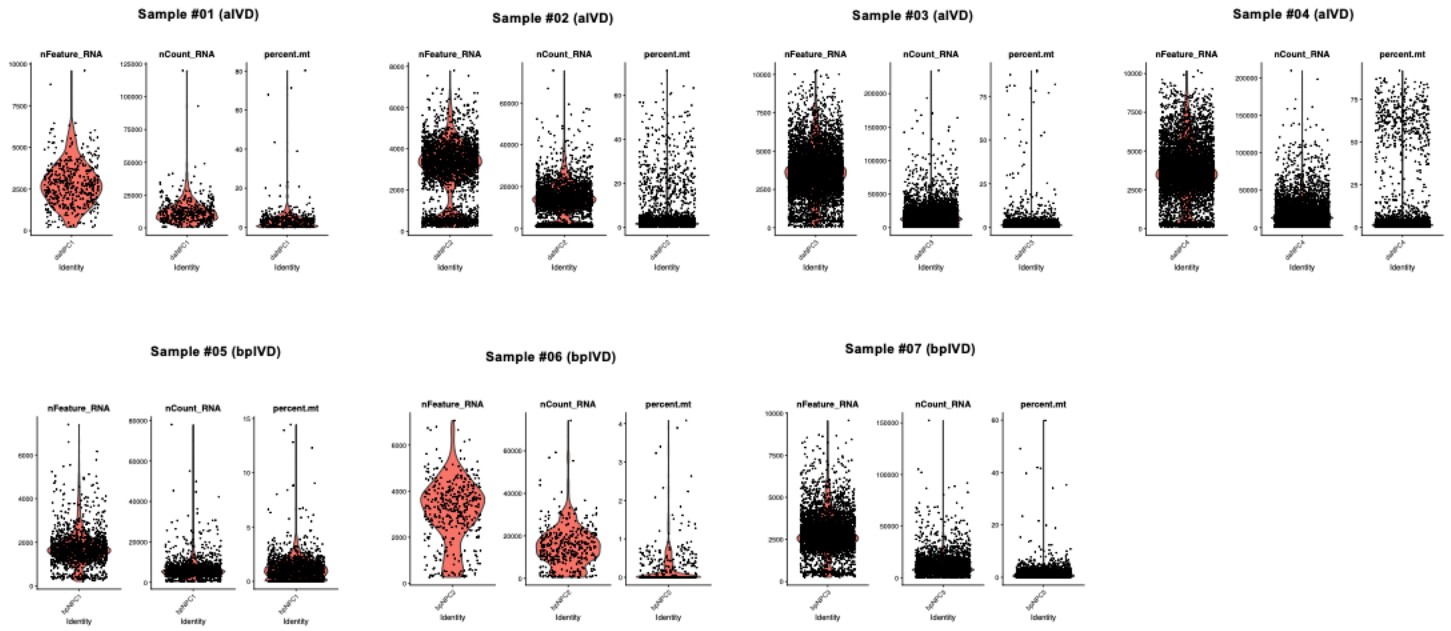


Figure S1. The number of genes detected per cell (nFeature_RNA), reads per gene (nCount_RNA), and the percent of mitochondrial genes (percent.mt) of single-cell data for each sample.

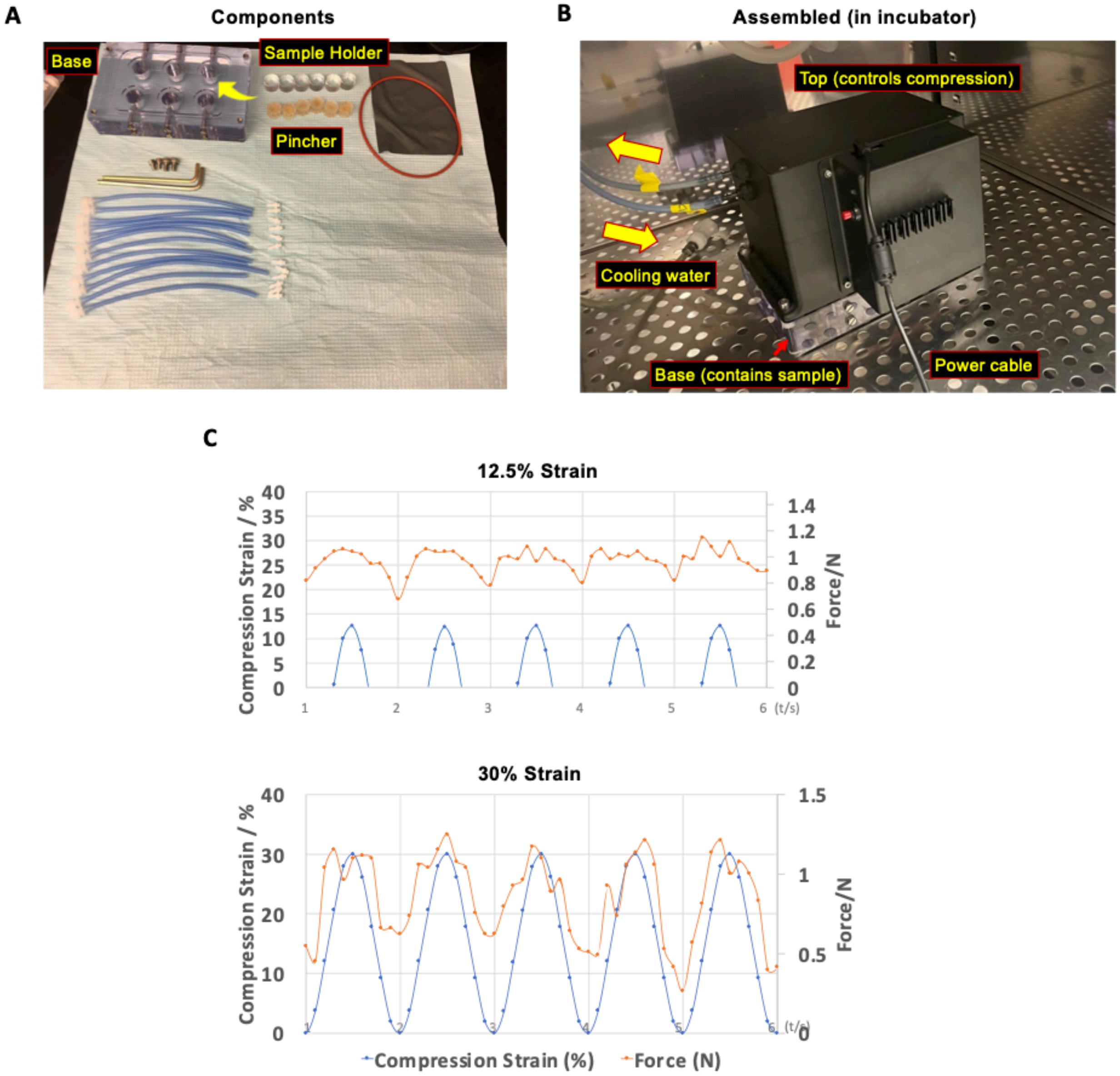


Figure S2. CellScale™ bioreactor. Optical photos of (A) the components for the CellScale™ bioreactor before assembly and (B) after assembled and placed in an incubator. The samples were placed in the sample holders which were placed in the wells on the base. (C) the compression strain and sensed force applied onto hydrogel construct loaded with NPCs. The frequency is 1Hz, and the strain is 12.5% and 30% respectively.

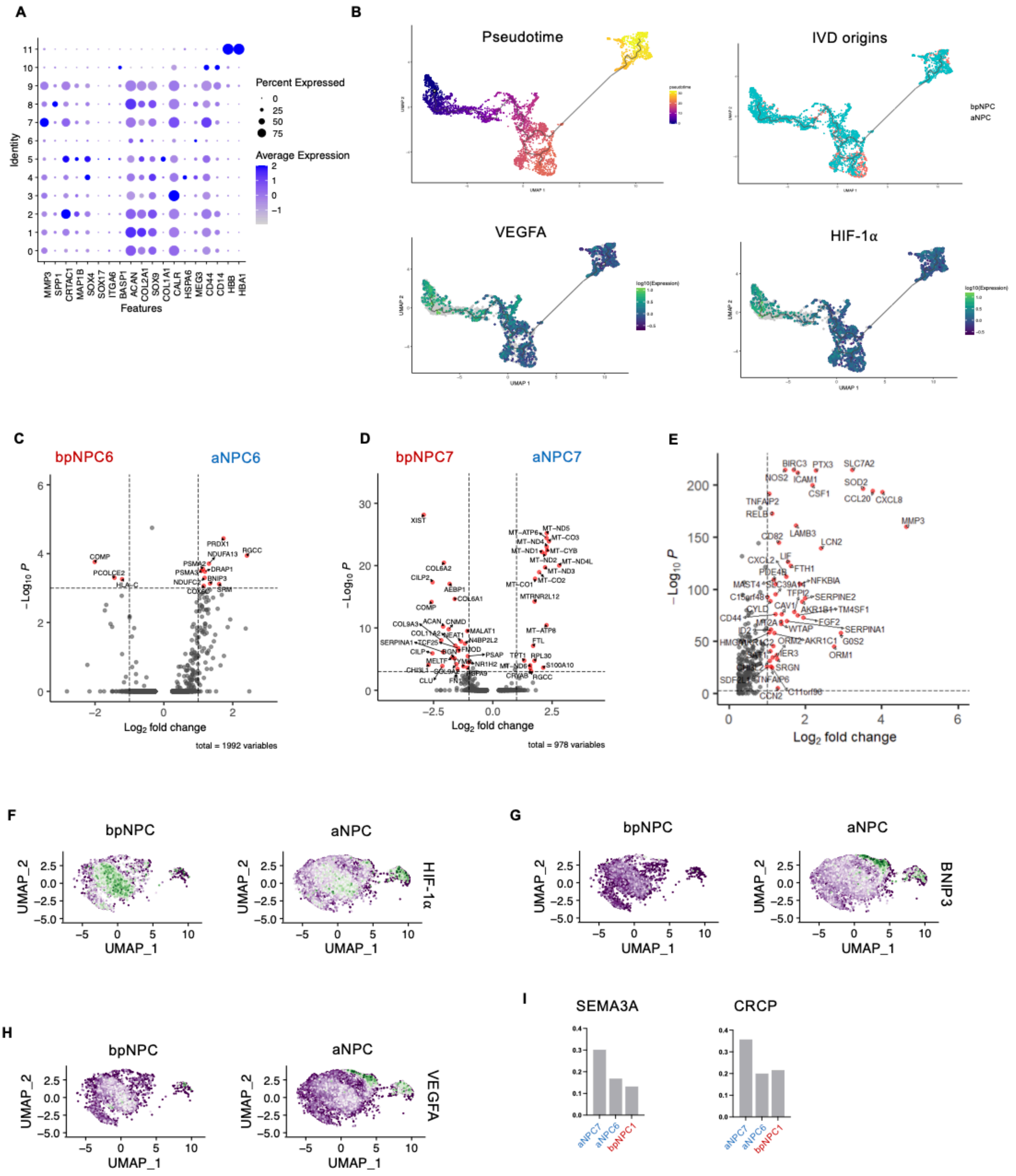


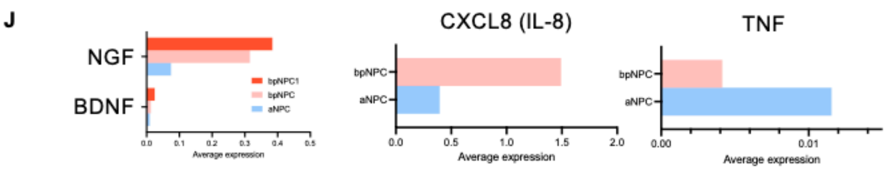


Figure S3. Supplemental figures for single-cell analyses of human IVD cells. **A)** The marker for annotating cell identifies for unsupervised clusters in human IVDs. Cluster 0-11 was annotated as NPC3, NPC4, FC, AFC1, NPC5, AFC2, NPC7, NPC1, NPC2, NPC6, IC, and RBC respectively. **B)** Developmental trajectory re-colored with pseudotime, IVD origns, expression level of VEGFA, or expression level of HIF-1α. Highly expressed markers differentiating **C)** aNPC6 and **D)** aNPC7 subtypes. **E)** Highly expressed markers for bpNPC1 cells (compared to other bpNPCs). Feature plots show expression levels of **F)** HIF-1α, **G)** BNIP3, and **H)** VEGFA in bpNPCs and aNPCs. **I)** Expression levels of SEMA3A and CRCP in aNPC7, aNPC6, and bpNPC1. J) Expression levels of NGF, BDNF in bpNPC1 cells, bpNPCs, and aNPCs and expression levels of IL-8, and TNF in bpNPCs and aNPCs.

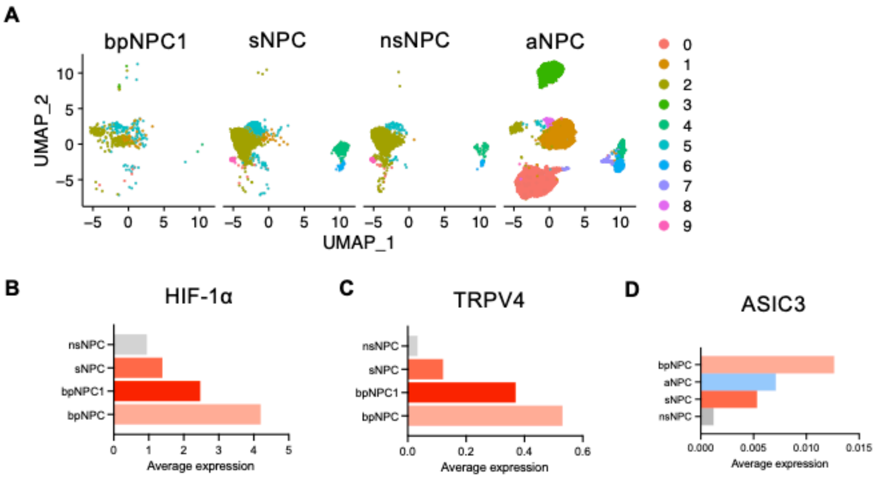


Figure S4. Supplemental figures for single-cell analyses of human nsNPC, sNPC, bpNPC, and aNPC cells. **A)** UMAP integrating bpNPC1, sNPC, nsNPC, and aNPC shows the aNPCs were very different after cryopreservation, thawing, and in vitro culture process (nsNPC). Gene expression levels of **B)** HIF-1α, **C)** TRPV4, and **C)** ASIC3

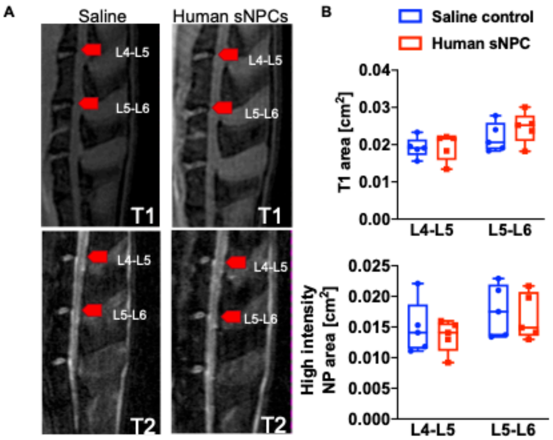


Figure S5. The μMRI images showing rat IVDs injected with sNPCs or saline. A) μMRI images of IVDs in sNPC-injected or saline-injected group at T1 and T2 directions. Red arrows point to the disc levels performed with intradiscal injections. B) The quantitative results for T1 area and high-density NP area for L4-L5 disc and L5-L6 disc (n=5). The μMRI images showed no evidence of IVD degeneration for IVDs injected with sNPCs or saline from T1 or T2 direction (Fig. 6D). No statistically difference of T1 area and high density NP area between the sNPC-injected and saline-injected group was detected in Fig. 6D. Our results showed that the intradiscal injection of sNPCs did not reduce the NP area.
